# Cortical Representations of Concrete and Abstract Concepts in Language Combine Visual and Linguistic Representations

**DOI:** 10.1101/2021.05.19.444701

**Authors:** Jerry Tang, Amanda LeBel, Alexander G. Huth

## Abstract

The human semantic system stores knowledge acquired through both perception and language. To study how semantic representations in cortex integrate perceptual and linguistic information, we created semantic word embedding spaces that combine models of visual and linguistic processing. We then used these visually-grounded semantic spaces to fit voxelwise encoding models to fMRI data collected while subjects listened to hours of narrative stories. We found that cortical regions near the visual system represent concepts by combining visual and linguistic information, while regions near the language system represent concepts using mostly linguistic information. Assessing individual representations near visual cortex, we found that more concrete concepts contain more visual information, while even abstract concepts contain some amount of visual information from associated concrete concepts. Finally we found that these visual grounding effects are localized near visual cortex, suggesting that semantic representations specifically reflect the modality of adjacent perceptual systems. Our results provide a computational account of how visual and linguistic information are combined to represent concrete and abstract concepts across cortex.

## Introduction

Humans learn about the world through both perception and language. The acquired knowledge is stored in cerebral cortex as semantic concept representations, which support a range of cognitive processes including language understanding. Many previous fMRI studies have found that concepts are represented near the perceptual systems through which they are commonly experienced (Binder and Desai, 2011; Harpaintner et al., 2020; Martin, 2016). These studies support grounded cognition theories, which hold that a concept’s semantic representation is formed through generalization or re-enactment of perceptual representations involved in learning the concept (Barsalou, 2008; Binder and Desai, 2011). Other studies have found that BOLD responses to words (Mitchell et al., 2008) and narratives (Huth et al., 2016; Wehbe et al., 2014) can be predicted using distributional word embeddings, which capture word co-occurrence statistics in language data. Distributional word embeddings lack explicit connections to the physical world (Bruni et al., 2014; Harnad, 1990), so their success in modeling brain responses demonstrates that semantic representations reflect word associations that can be learned from language alone. Together these findings suggest that semantic representations contain both perceptual and linguistic information (Andrews et al., 2014). However, little is known about how these different sources of information are combined to form semantic representations in each cortical region.

One open question is whether different cortical regions represent concepts using different amounts of perceptual and linguistic information. Grounded cognition theories predict that representations in each semantically selective cortical region reflect how information is represented in adjacent perceptual systems (Barsalou, 2008; Binder and Desai, 2011). For instance, these theories predict that cortical regions near the visual system represent concepts using visual information. We might similarly expect cortical regions near the language system to represent concepts using information about language usage, such as distributional word co-occurrence. However, there is little work directly assessing these theories by comparing semantic representations in each cortical region to computational models of perceptual and linguistic processing (Anderson et al., 2019). A second open question is whether concrete and abstract concepts are represented using different amounts of perceptual and linguistic information. Previous studies (Binder et al., 2005; Paivio, 1991) suggest that concrete concepts—which are directly experienced through perception—contain more perceptual information, but this relationship has not been directly tested using fMRI. Furthermore, the role of perceptual information in representing abstract concepts—which are not directly experienced through perception—is under debate. Traditional views hold that abstract concepts are represented solely by linguistic information (Dove, 2009; Paivio, 1991), while recent studies suggest that abstract concepts contain some amount of perceptual information (Harpaintner et al., 2018). A third open question is how the semantic system represents concepts experienced through multiple perceptual modalities. Grounded cognition theories predict that concepts are represented near each perceptual system through which they are experienced, in a format that specifically reflects that perceptual modality (Barsalou, 2008; Martin, 2016). For instance, visual features of “hammer” might be represented near visual cortex, while tactile features of “hammer” might be represented near somatosensory cortex. Alternatively, concepts could be represented across cortex in a format that integrates information from multiple different perceptual modalities. For instance, each cortical region selective for “hammer” might simultaneously represent its visual, tactile and auditory features.

Here, we investigated these questions by constructing a computational model of how *visual* and *linguistic* information combine to form semantic representations. We first modeled visual and linguistic representations as separate word embedding spaces. Embedding spaces represent each word using a high-dimensional vector, and quantify the similarity between each pair of words using the dot product between their corresponding vectors. Since our subjects have learned about concepts through both vision and language, we next modeled each word’s semantic representation by concatenating its visual and linguistic embeddings, making the semantic similarity between each pair of words a combination of their visual and linguistic similarities. Because the relative amount of visual and linguistic information may differ across brain regions or concepts, we weighted the visual and linguistic embeddings for each word prior to concatenation. By varying the weights on the visual and linguistic embeddings, we were able to construct a spectrum of semantic spaces that can capture different possibilities for how each word’s semantic representation combines its visual and linguistic representations.

We compared the different semantic embedding spaces to concept representations in each cortical region using a natural language fMRI experiment. In this experiment, BOLD fMRI responses were collected from seven human subjects as they listened to over five hours of narrative stories from *The Moth Radio Hour* (**Figure 1A**). These stories activate the semantic representations of thousands of concepts common in daily life. We then fit voxelwise encoding models that separately predict the fMRI data in each subject from the stimulus words (Huth et al., 2016; Jain and Huth, 2018; Wehbe et al., 2014). An encoding model uses regularized linear regression to estimate a set of weights for each voxel that predict how each word influences BOLD responses in that voxel. Encoding models were fit using an embedding space prior, which enforces that similar words in the embedding space should have similar encoding weights (Nunez-Elizalde et al., 2019). Since successful models of the brain should be able to generalize to new natural stimuli (Hamilton and Huth, 2018), encoding models were evaluated by predicting BOLD responses to stories that were not used for model estimation, and then computing the correlation between predicted and actual responses (**Figure 1B**).

**Figure 1.**
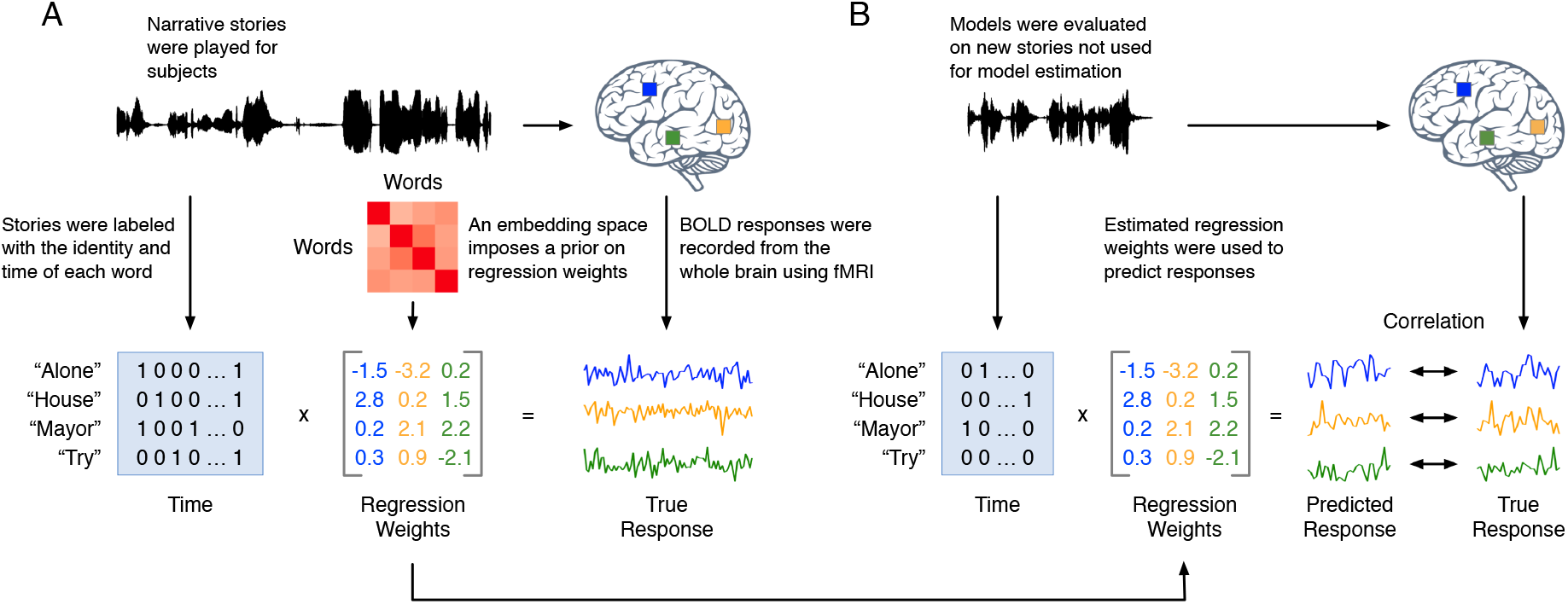
Natural language fMRI experiment. (**A**) Seven human subjects listened to over 5 hours of narrative stories while BOLD responses were measured using fMRI. A stimulus matrix was constructed by identifying the words spoken at each point in time in the stories. A regularized, linearized finite impulse response regression model was then estimated for each cortical voxel using a word embedding space prior. The estimated encoding model weights describe how words in the stories influence BOLD signals in each cortical voxel. The prior enforces that similar words in the embedding space should have similar encoding model weights. (**B**) Models were tested on stories that were not included in the model estimation procedure. Generalization performance for a test story was computed as the linear correlation between the predicted BOLD responses to the test story and the observed BOLD responses.

To quantify how much visual or linguistic information is represented in each cortical region, we fit separate voxelwise encoding models using embedding spaces that range from fully linguistic to fully visual. In voxelwise modelling, the embedding space that best reflects a voxel’s semantic representations will yield the best generalization performance. We thus operationalized the *representational format* of each voxel as the semantic embedding space with the best generalization performance.

## Results

### Construction of visual, linguistic, and semantic embedding spaces

In order to assess the amount of visual and linguistic information that is incorporated into semantic representations, we first needed to construct computational models of visual and linguistic processing. We did that here using separate visual and linguistic word embedding spaces, which are then combined in different ratios to create semantic embedding spaces.

We modeled linguistic representations using distributional word embeddings, which assign each word a vector based on its co-occurrence statistics with a set of target words across a large corpus. Such embeddings have been shown to capture meaningful linguistic associations (Deerwester et al., 1990; Lund and Burgess, 1996), and are widely used as computational models of lexical semantics (Pennington et al., 2014). Here, we used a distributional embedding space previously shown to model BOLD responses to narrative stories (de Heer et al., 2017; Deniz et al., 2019; Huth et al., 2016). While co-occurrence statistics may implicitly capture some degree of perceptual similarity (Riordan and Jones, 2011), they do not incorporate explicit information about the physical world (Glenberg and Robertson, 2000; Harnad, 1990), making them an appropriate model of knowledge acquired through language. Words that occur in similar linguistic contexts will have similar linguistic embeddings, and will thus be considered linguistically similar.

We modeled visual representations using image embeddings extracted from convolutional neural networks (CNNs). We first defined a diverse pool of *visual words*, which refer to entities or events that can be experienced through vision (see **Methods** for details). For each visual word, we sampled 100 related natural images from ImageNet (Deng et al., 2009). Recent studies (Cadieu et al., 2014; Eickenberg et al., 2017; Güçlü and van Gerven, 2015; Khaligh-Razavi and Kriegeskorte, 2014; Yamins et al., 2014) have shown that primate visual processing is well-modeled by CNNs trained to identify objects in images (Chatfield et al., 2014; Krizhevsky et al., 2012; Sermanet et al., 2013; Zeiler and Fergus, 2014). We used a similar CNN (VGG16; Simonyan and Zisserman, 2015) to extract embedding vectors for each image. The visual embedding for each visual word was then obtained by averaging the extracted CNN embeddings across the 100 sampled images. Words with referents that evoke similar responses in visual cortex will have similar visual embeddings, and will thus be considered visually similar.

We next estimated visual embeddings for *non-visual words*. While non-visual words refer to concepts that cannot be directly experienced through vision, recent studies suggest that their representations may nonetheless contain some amount of visual information (Harpaintner et al., 2018). To capture this, we developed a *perceptual propagation* method that represents non-visual words by combining the visual embeddings of linguistically associated visual words (similar to Collell et al., 2017). For each non-visual word *w*, we fit a linear regression θ_*w*_ to reconstruct its linguistic embedding as a weighted sum of the linguistic embeddings of visual words. Visual words that are linguistically associated with *w* will have high weights in θ_*w*_. We then predicted a visual embedding for *w* by applying the same linear weights θ_*w*_ to the visual embeddings of the visual words. Non-visual words will thus be considered visually similar if they are linguistically associated with visually similar words. For instance, the non-visual words “famous” and “lonely” are dissimilar in the linguistic embedding space but similar in the visual embedding space, as they are respectively associated with the visually similar words “musician” and “friend”. **Figure 2A** summarizes the process of creating visual and linguistic embedding spaces.

**Figure 2.**
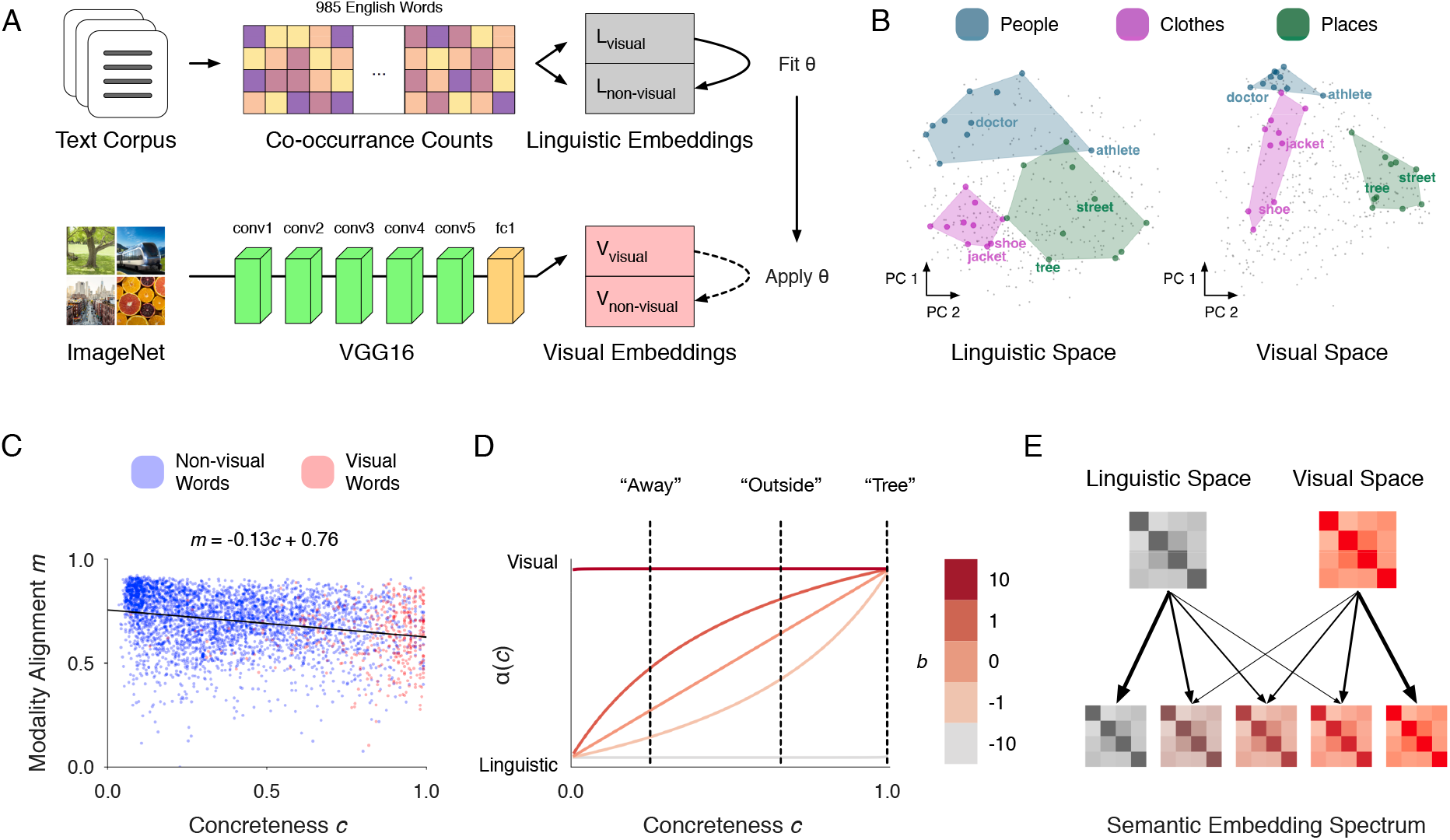
Construction of visual, linguistic, and semantic embedding spaces. (**A**) Linguistic embedding vectors were constructed from distributional co-occurrence statistics in a large external corpus. Visual embedding vectors for visual words were constructed by sampling 100 images from ImageNet for each word and averaging embeddings extracted from a VGG16 convolutional neural network. Visual embedding vectors for non-visual words were constructed using a perceptual propagation method θ that represents each non-visual word as a linear combination of the visual embeddings of associated visual words (see **Methods** for details). (**B**) The visual and linguistic embedding spaces were visualized by projecting the embedding of each visual word onto the first two principal components of the embedding space. The visual and linguistic embedding spaces structure words into similar high-level *people, clothing*, and *place* categories. However, fine-grained similarities within each category differ across embedding spaces. Words with visually similar referents (e.g. *people*) are more similar in the visual space, while words that occur in similar linguistic contexts (e.g. *clothes*) are more similar in the linguistic space. (**C**) For each word, a *modality alignment score*— computed as the linear correlation between its visual similarities and linguistic similarities with other words—was plotted against a *concreteness score* derived from behavioral judgments. Visual words were colored red, and non-visual words blue. Modality alignment scores are weakly anticorrelated with concreteness scores, suggesting that visual and linguistic embedding spaces differ more for concrete words than for abstract words. Nonetheless, visual and linguistic similarity differ to some degree even for highly abstract words, demonstrating that the visual embedding space represents abstract words using visual information absent from the linguistic embedding space. (**D**) Semantic embedding spaces were constructed by concatenating visual and linguistic embeddings for each word. Prior to concatenation, the visual and linguistic embeddings were weighted by a function α_concrete_ of each word’s concreteness score, and the total amount of visual information for each word was controlled by a parameter *b*. Varying *b* creates a semantic embedding spectrum that interpolates between the linguistic embedding space and the visual embedding space. Intermediate spaces in the semantic embedding spectrum represent each word as a combination of visual and linguistic information.

Before using the visual and linguistic embedding spaces to model semantic representations in the brain, we first tested whether they capture different notions of similarity. We did this by defining semantic categories consisting of *people, clothing*, and *place* words and then identifying qualitative differences in how these categories are represented across embedding spaces (**Figure 2B**). We visualized each embedding space by using principal components analysis (PCA) to project the embedding of each visual word onto two dimensions. PCA projects words with similar embeddings to nearby points in 2D space, and those with very different embeddings to distant points. First, we found that both embedding spaces contain distinct *people, clothing*, and *place* clusters, reflecting previous findings that visual and linguistic embedding spaces structure concepts into similar categories (Riordan and Jones, 2011). However, we found that relationships within each category differed between the visual and linguistic embedding spaces. For instance, *people* words (such as “doctor”, “athlete”, and “friend”) are close together in the visual space, reflecting their shared visual features, and far apart in the linguistic space, reflecting their diverse linguistic contexts. In contrast, *clothing* words (such as “jacket”, “shoe”, and “hat”) are far apart in the visual embedding space, reflecting their diverse visual features, and close together in the linguistic embedding space, reflecting their shared linguistic contexts. This qualitative analysis suggests that the visual and linguistic embedding spaces structure concepts into similar high-level categories, but capture fine-grained notions of visual and linguistic similarity within each category.

While the previous analysis shows that visual and linguistic embedding spaces differ within visual categories like *people* and *clothing*, it is unclear whether they also differ for more abstract words. Our perceptual propagation method predicts that non-visual words (which tend to be more abstract) acquire visual information through associations with visual words. However, for highly abstract words that are not strongly associated with any visual words, the estimated visual embeddings may not contain any meaningful visual information. In that case, we might expect no difference between the visual and linguistic embedding spaces. To test this possibility, we quantified the difference between visual and linguistic model representations for each individual word. We did this by constructing visual and linguistic *similarity vectors* for each word that contain its visual and linguistic similarity with every other word. We then computed a *modality alignment score* for each word as the linear correlation between its visual and linguistic similarity vectors. We plotted each word’s modality alignment score against a *concreteness score* derived from a separate dataset of behavioral judgments about word concreteness (Brysbaert et al., 2014; see **Methods**). We found that modality alignment scores are anticorrelated with concreteness scores (*r* = -0.26), suggesting that the visual and linguistic embedding spaces differ more for concrete words than for abstract words. Nonetheless, we found that the visual and linguistic embedding spaces differ to some degree even for highly abstract words, suggesting that the visual embedding space represents abstract words using some visual information that is absent from the linguistic embedding space (**Figure 2C**).

Finally, we combined the visual and linguistic embedding spaces into semantic embedding spaces to model how concepts are represented in the brain’s semantic system. Since our subjects have learned about the world through both vision and language, we expect each word’s semantic representation to combine the two information sources. Semantic embedding spaces formalize this hypothesis by representing each word as a concatenation of its visual and linguistic embeddings. Since different words may contain different amounts of visual and linguistic information, each word *w* is assigned a *modality weight* α_*w*_ such that its visual embedding is weighted by α_*w*_ and its linguistic embedding is weighted by (1 - α_*w*_) prior to concatenation. The semantic similarity between each pair of words is thus modeled as a combination of their visual and linguistic similarities, weighted by the modality weights of both words (see **Methods**). Under this model, each semantic embedding space is generated by a vector **α** of modality weights across the words, and captures a different possibility for how visual and linguistic information are combined to represent each word. For example, setting α = 1 for all words would capture the hypothesis that all concepts are represented in a visual format, while setting α = 1 for concrete words and α = 0 for abstract words would capture the hypothesis that only concrete concepts are represented in a visual format.

The space of **α** vectors—and thus the number of possible semantic embedding spaces—is infinitely large. To constrain this space, we only considered modality weights that are monotonically increasing functions α_concrete_ (see **Methods**) of concreteness score *c*. This hypothesis reflects previous findings that more concrete words appear to contain more perceptual information (Harpaintner et al. 2018, Anderson et al. 2019). The α_concrete_ model has a single parameter *b* that biases the degree to which each word is represented by visual information (**Figure 2D**). When *b* is small, α_concrete_(*c*) approaches 0 for all values of *c*, causing all words to be represented solely by their linguistic embeddings. As *b* increases, more concrete words are represented by more visual information. When *b* is large, α_concrete_(*c*) approaches 1 for all values of *c*, causing all words to be represented solely by their visual embeddings. We tested a range of *b* values (−10, -1, 0, 1, 10) that induce semantic embedding spaces ranging from *fully linguistic* (*b* = -10) to *fully visual* (*b* = 10). We considered all embedding spaces containing some amount of visual information (*b* = -1, 0, 1, 10) to be *visually grounded*. This semantic embedding spectrum captures a diverse set of hypotheses for how visual and linguistic information are combined in each word’s semantic representation (**Figure 2E**).

### Representational format of cortical regions near visual and language systems

We first compared semantic embedding spaces to characterize the representational format of each semantically selective cortical region. Grounded cognition theories (Barsalou, 2008; Binder and Desai, 2011) predict that cortical regions near the visual system respond similarly to visually similar words, and should thus be best modeled by visually grounded embedding spaces. Conversely, we predict that cortical regions near the language system respond similarly to linguistically similar words, and should thus be best modeled by the fully linguistic embedding space. Previous studies have tested whether cortical regions are better modeled by an experiential embedding space, a linguistic embedding space, or a multimodal embedding space that combines the two information sources (Anderson et al., 2019). However, this experiential embedding space reflects coarse-grained behavioral ratings of whether concepts are experienced through similar perceptual modalities (such as whether each concept “has a characteristic or defining color”), rather than fine-grained similarity within a specific perceptual modality. Furthermore, multimodal embeddings were modeled in (Anderson et al., 2019) as unweighted concatenations of perceptual and linguistic embeddings, which implicitly assumes that each concept is represented by the same amount of perceptual and linguistic information. Our semantic embedding spectrum differs from these previous models in two important ways: CNN embeddings explicitly reflect fine-grained visual similarity (Eickenberg et al., 2017), and different semantic embedding spaces model different hypotheses for how each concept’s semantic representation combines visual and linguistic information.

For each subject, we fit voxelwise encoding models using each space in the semantic embedding spectrum, and then tested the generalization performance of each model on held-out data. We identified *semantic system voxels* that were significantly predicted under any space in the embedding spectrum (q(FDR) < 0.05, blockwise permutation test; see **Methods**). Our encoding models significantly predicted up to 18 percent of cortical voxels in each subject. These semantic system voxels were located in broad regions of prefrontal cortex, temporal cortex, and parietal cortex (see **Figure S1** for encoding model performance across cortex) that align with semantically selective regions reported in previous studies (Binder et al., 2009; Huth et al., 2016).

To compare the different semantic spaces, we aggregated model performance across semantic system voxels near known vision and language regions of interest (ROIs), which were identified in each subject using separate localizer data (see **Methods** for details). For vision ROIs we defined the fusiform face area (FFA), parahippocampal place area (PPA), occipital place area (OPA), retrosplenial cortex (RSC), and extrastriate body area (EBA). For language ROIs we defined the auditory cortex (AC), Broca’s area, and superior premotor ventral speech area (sPMv). The performance of each embedding space around each ROI was first summarized by averaging encoding model generalization performance across all semantic system voxels within 15mm of the ROI along the cortical surface. We then defined the *visual grounding score* for each visually grounded space around an ROI as the difference between its encoding performance and that of the fully linguistic space (**Figure 3**). If any visually grounded spaces have a positive visual grounding score around an ROI, it would suggest that semantically selective cortical regions near the ROI tend to represent concepts using some amount of visual information. If all visually grounded spaces have a negative visual grounding score, it would suggest that semantically selective cortical regions near the ROI tend to represent concepts using mostly linguistic information.

**Figure 3.**
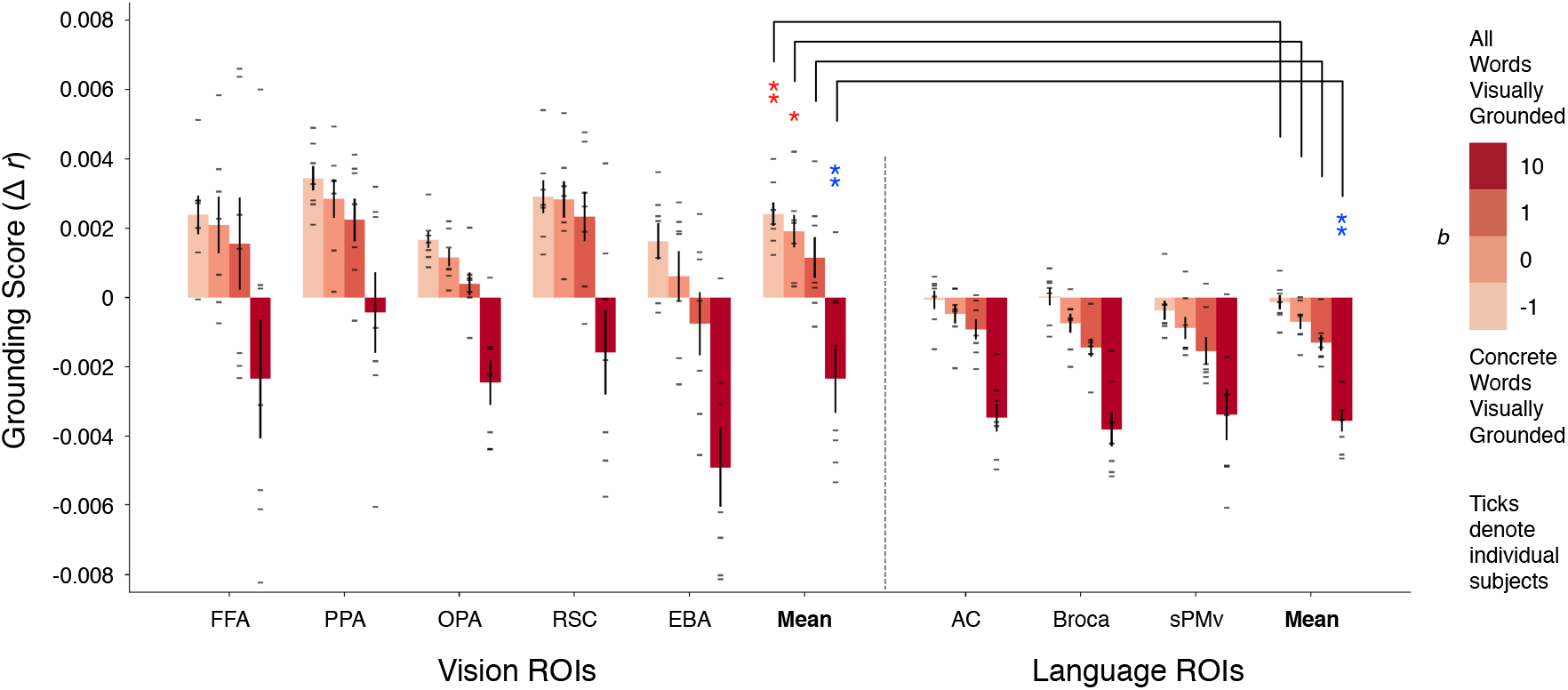
Representational format of cortical regions near visual and language systems. Encoding models were fit using each space in a semantic embedding spectrum ranging from fully linguistic to fully visual. Vision- and language-related functional regions of interest (ROIs) were identified for each subject using separate localizer data. Embedding space performance around an ROI was quantified by averaging encoding model generalization performance (linear correlation *r*) across all significantly-predicted voxels within 15 mm of the ROI along the cortical surface. For each visually grounded embedding space, *visual grounding score*—defined as the performance improvement over the fully linguistic embedding space—was averaged across subjects and plotted for each ROI and ROI type (vision and language). Ticks denote visual grounding scores for individual subjects. Error bars indicate the standard error of the mean across subjects (*n* = 7). We used a linear mixed-effects model to compare visual grounding score around vision and language ROIs for each visually grounded embedding space. Significance was tested for each ROI type (vision, language); red asterisks indicate that a visually grounded space performs significantly better than the fully linguistic space, and blue asterisks indicate that a visually grounded space performs significantly worse (*, q(FDR) < 0.05; **, q(FDR) < 10^−2^; ***, q(FDR) < 10^−3^, ****, q(FDR) < 10^−4^). Brackets signify that the visual grounding score of each visually grounded space is significantly higher around vision ROIs than around language ROIs (q(FDR) < 0.05). These results show that visually grounded embedding spaces significantly outperform the fully linguistic embedding space near vision ROIs, but not language ROIs.

We used a linear mixed-effects model to compare visual grounding score for each visually grounded space (4 levels) across ROI type (2 levels: vision, language) with ROI identity as a random effect nested in subject identity. This test showed that visual grounding score varies significantly across embedding spaces (Wald χ^2^ test, *p* < 10^−4^) and ROI type (*p* < 10^−4^). There was also a significant interaction between embedding space and ROI type (*p* = 0.012), demonstrating that semantic embedding spaces have different patterns of generalization performance across vision and language ROIs. A post hoc test comparing the visual grounding score of each visually grounded space against the null hypothesis of zero found that multiple visually grounded spaces (*b* = -1, 0) significantly outperformed the fully linguistic space around vision ROIs (q(FDR) < 0.05), while no visually grounded spaces significantly outperformed the fully linguistic space around language ROIs. A post hoc test comparing visual grounding score around vision and language ROIs found that every visually grounded space had a significantly higher visual grounding score around vision ROIs than around language ROIs (q(FDR) < 0.05).

**Figure 3** shows these differences between the semantic embedding spaces around visual and language ROIs. The small size of these effects is likely a consequence of our encoding framework and the large amount of fMRI data (5 hours per participant) that was used. In a regularized encoding model, different embedding spaces impose different priors on the model weights (Nunez-Elizalde et al., 2019), but as the amount of training data increases, the model can learn accurate weights from the data alone. Comparing embedding spaces by fitting encoding models on large fMRI datasets thus reveals small but significant differences in performance.

Our results provide fMRI evidence that cortical regions near the visual system represent concepts using both visual and linguistic information, while cortical regions near the language system represent concepts using mostly linguistic information (Barsalou, 2008; Binder and Desai, 2011). These results are markedly different from previous fMRI studies, which found that multimodal embedding spaces outperform linguistic embedding spaces in superior temporal and inferior frontal regions, but not in cortical regions near the visual system (Anderson et al., 2019). The success of our visually grounded embedding spaces in these latter regions suggests that semantic representations near the visual system specifically reflect fine-grained visual information, which is captured in our CNN embeddings but not in previous experiential embeddings.

### Visual grounding of concrete and abstract concepts near visual cortex

The previous analyses show that concept representations in regions near visual cortex are better modeled by visually grounded embedding spaces that combine visual and linguistic information (*b* = -1, 0, 1) than by embedding spaces that solely reflect linguistic (*b* = -10) or visual (*b* = 10) information. In these intermediate visually grounded embedding spaces, the relative weighting of each word’s visual and linguistic embeddings was selected to be a function α_concrete_ of the word’s concreteness score. The α_concrete_ model captures two major hypotheses for how semantic representations combine visual and linguistic information. First, α_concrete_ is a monotonically increasing function of concreteness. This models the hypothesis that more concrete concepts are represented by more visual information while more abstract concepts are represented by more linguistic information (Paivio, 1991). Second, the visually grounded parameterizations of α_concrete_ (*b* = -1, 0, 1, 10) assign a positive weight to every word, meaning that even abstract words are represented to some extent by their estimated visual embeddings. This models the hypothesis that abstract concepts are represented using some amount of perceptual information from linguistically associated concrete concepts. In the following analyses we focused on semantically selective regions near visual cortex, and directly tested these two hypotheses by comparing the α_concrete_ model against alternative modality weight models.

To quantify how well a modality weight model explains semantic representations near visual cortex, we fit an encoding model using the semantic embedding space that it generates. We then averaged encoding model performance (linear correlation *r*) across semantic system voxels within 15mm of vision ROIs. Before comparing against alternative modality weight models, we selected the best visually grounded α_concrete_ model across the tested voxels (*b* = -1) using separate validation data (see **Methods**).

Previous theories have proposed that concrete concept representations contain more perceptual information, while abstract concept representations contain more linguistic information (Paivio, 1991). However, this hypothesis has not been directly tested at the level of individual words using fMRI. Here, we conducted a permutation test to quantify whether the concreteness of each concept explains the amount of visual and linguistic information in that concept’s representation. We conducted 1,000 trials in which we permuted concreteness scores across words before computing modality weights under the α_concrete_ model. Each trial *t* produced a vector of modality weights **α**_*t*_ corresponding to a different permutation of the concreteness-derived modality weights **α**_concrete_ (**Figure 4A**). We then evaluated encoding model performance under the semantic embedding space generated by **α**_*t*_. If the amount of visual and linguistic information in each concept representation does not reflect concreteness, then model performance using the true concreteness scores should not be substantially different from performance using randomly permuted concreteness scores. However, if the amount of visual and linguistic information in each concept representation can be explained by concreteness, then model performance using the true concreteness scores should be much higher than performance using randomly permuted concreteness scores.

**Figure 4.**
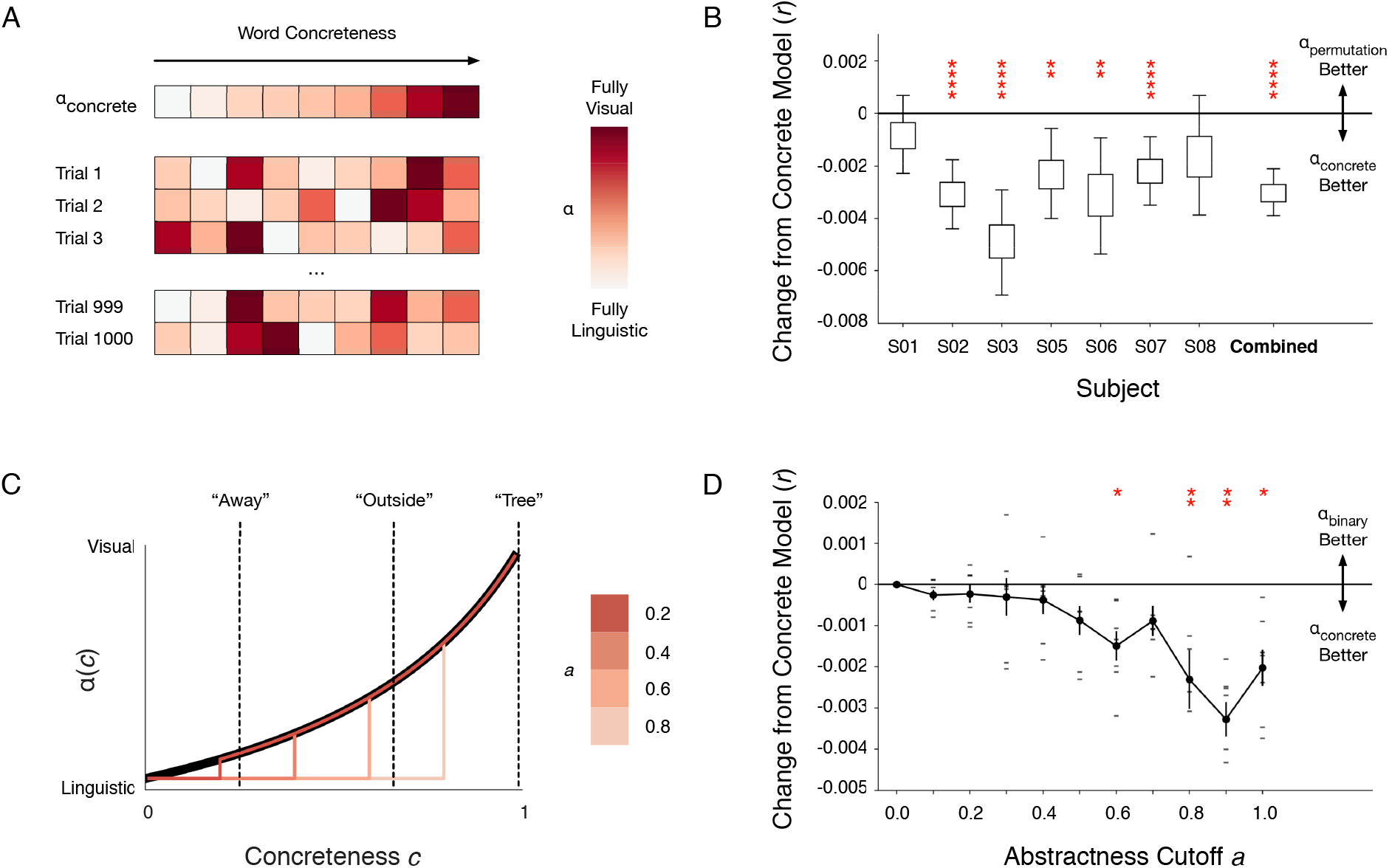
Visual grounding of concrete and abstract concepts near visual cortex. Encoding models fit under a visually grounded α_concrete_ modality weight model were compared to encoding models fit under alternative modality weight models. Performance for each encoding model was quantified by averaging generalization performance (linear correlation *r*) across all significantly-predicted voxels within 15 mm of vision ROIs along the cortical surface. **(A)** A permutation test was performed to quantify whether concreteness explains the amount of visual and linguistic information in each concept representation. In each trial, concreteness scores were permuted across words before modality weights were computed under the α_concrete_ model. **(B)** The difference between the permutation distribution of encoding performance and the observed encoding performance of the α_concrete_ model was first plotted for each subject, and then aggregated across the seven subjects. Boxes indicate the interquartile range of the differences; whiskers indicate the 2.5th and 97.5th percentiles. If the true amount of visual information in each concept representation increases with concreteness, the permutation distribution should be lower than the observed test statistic. If the true amount of visual information in each concept representation is not related to concreteness, the permutation distribution should not, on average, differ from the observed test statistic. Red asterisks signify that the permutation distribution is significantly lower than the α_concrete_ model performance for five of seven individual subjects, and combined across subjects. (*, q(FDR) < 0.05; **, q(FDR) < 10^−2^; ***, q(FDR) < 10^−3^, ****, q(FDR) < 10^−4^). **(C)** The α_binary_ model modifies the α_concrete_ model to assign a modality weight of 0 to all words with concreteness scores below an abstractness cutoff. Abstractness cutoffs operationalize the hypothesis that certain abstract concepts are represented solely by linguistic information. **(D)** Model performance under the α_binary_ model for different abstractness cutoffs was compared to model performance under the α_concrete_ model. Error bars indicate the standard error of the mean across (*n* = 7) subjects. Red asterisks signify that an α_binary_ model performed significantly worse than the α_concrete_ model (*, q(FDR) < 0.05; **, q(FDR) < 10^−2^; ***, q(FDR) < 10^−3^, ****, q(FDR) < 10^−4^). These results suggest that many abstract concept representations (*c* < 0.6) near visual cortex contain some amount of visual information.

We found that the encoding performance of the α_concrete_ model was significantly higher than the permutation distribution of encoding performance when combined across subjects (q(FDR) < 10^− 4^), and individually for 5 of 7 subjects (q(FDR) < 10^−2^) (**Figure 4B**). These results suggest that the amount of visual and linguistic information in each concept representation is significantly related to concreteness; more concrete concepts contain more visual information, while more abstract concepts contain more linguistic information.

We next addressed the question of whether abstract concept representations contain *any* perceptual information. Traditional views propose a binary in which concrete concepts are represented by perceptual and linguistic information, while abstract concepts are represented solely by linguistic information (Dove, 2009; Paivio, 1991). Conversely, recent behavioral studies suggest that many abstract concepts contain some amount of perceptual information (Borghi et al., 2017; Harpaintner et al., 2020, 2018). Extending these recent findings, our perceptual propagation method estimates visual embeddings of non-visual words by combining the visual embeddings of linguistically associated visual words. The visually grounded α_concrete_ models (*b* = -1, 0, 1, 10) then assign each abstract word a positive weight on its estimated visual embedding, modeling the hypothesis that abstract concept representations contain visual information from linguistically associated visual concepts. Here, we directly tested if abstract concepts are better modeled by including some amount of this associated visual information, or solely by linguistic information.

We operationalized the traditional binary view of abstractness by defining *abstractness cutoffs* on concreteness scores. For each abstractness cutoff, words with concreteness scores below the cutoff value were represented solely by their linguistic embeddings, while words with concreteness scores above the cutoff were represented by a weighted concatenation of visual and linguistic embeddings. Formally, this binary view of abstractness is captured by a modality weight model α_binary_ with an abstractness cutoff parameter *a* (**Figure 4C**). α_binary_ maps concreteness scores *c* below the cutoff to 0 and maps concreteness scores above the cutoff to α_concrete_(*c*) (see **Methods**). If setting an abstractness cutoff increases performance relative to α_concrete_, it would suggest that words with concreteness scores below the cutoff tend to be represented solely by linguistic information. However, if setting an abstractness cutoff decreases performance relative to α_concrete_, it would suggest that words with concreteness scores below the cutoff tend to be represented by a combination of visual and linguistic information. We tested the α_binary_ model for a range of abstractness cutoffs (**Figure 4D**). We used a linear mixed-effects model to compare the performance difference between each α_binary_ model (11 levels) and the α_concrete_ model with subject identity as a random effect. This test showed that performance difference varies significantly across α_binary_ models (Wald χ^2^ test, *p* < 10^−4^). A post hoc test comparing the performance between each α_binary_ model and the α_concrete_ model found that α_binary_ models with abstractness cutoffs of 0.6, 0.8, 0.9, and 1.0 performed significantly worse than the α_concrete_ model (q(FDR) < 0.05). These results suggest that many abstract concepts (*c* < 0.6) are represented in a format that includes perceptual information from linguistically associated concrete concepts.

### Representational format of concrete concepts across cortex

Our results suggest that cortical regions near the visual system represent concepts in a format that explicitly reflects visual information (**Figure 4**), supporting theories that the semantic representations of concrete concepts are formed through reuse of representations in adjacent perceptual systems (Barsalou, 2008; Binder and Desai, 2011). However, concrete concepts tend to be experienced through multiple perceptual modalities, and not solely vision (Lynott et al., 2020). Thus it remains unclear how their semantic representations might combine information from different perceptual systems. Grounded cognition theories predict that concrete concepts are represented near each perceptual system through which they are experienced using information from that particular perceptual modality (Barsalou, 2008; Martin, 2016). Alternatively, concrete concepts could be represented across cortex in a common multimodal format that combines representations from multiple perceptual modalities. For instance, (Amedi et al., 2001) found that certain regions in lateral occipital cortex are activated when subjects either view or hold an object, suggesting that these regions contain multimodal representations of object shape.

Our results thus far are consistent with both possibilities. Voxels near visual cortex may be best modeled by visually grounded embedding spaces because their representations specifically reflect visual information. However, it may also be possible that all concrete concepts are represented in a multimodal format that includes some visual information as well as information from other perceptual systems. In this case, voxels near visual cortex may be best modeled by visually grounded embedding spaces simply because they represent concrete concepts. To differentiate these possibilities, we quantified the *concrete selectivity* and *visual grounding* of each voxel in the semantic system. If concrete concepts are represented near each perceptual system in a format that specifically reflects the corresponding modality, we would expect visually grounded embedding spaces to only perform well near visual cortex. However, if concrete concepts are represented in a common multimodal format across cortex, we would expect visually grounded embedding spaces to perform well in all cortical regions that represent concrete concepts.

We defined a *concrete selectivity score* for each voxel by projecting its encoding model weights onto the vector of concreteness scores for each word. Voxels which tend to respond more to concrete words than abstract words will have positive concrete selectivity scores, while voxels which tend to respond more to abstract words than concrete words will have negative concrete selectivity scores. We defined a *visual grounding score* for each voxel as the difference in encoding model performance between the best performing visually grounded embedding space across cortex (*b* = -1; see **Methods**) and the fully linguistic embedding space. Voxels that represent concepts using some amount of visual information will have positive visual grounding scores, while voxels which represent concepts using mostly linguistic information will have negative visual grounding scores.

We projected the concrete selectivity and visual grounding scores for each semantic system voxel onto a cortical flatmap. Each voxel was assigned a brightness based on its concrete selectivity score and a color based on its visual grounding score. In this visualization, concrete selective voxels appear red if they are best modeled by the visually grounded space, and blue if they are best modeled by the linguistic space. Abstract selective voxels appear black. The resulting map (**Figure 5A**; see **Figure S1** for other subjects) shows that voxels near perceptual systems (specifically visual cortex, somatosensory cortex, and auditory cortex) tend to be concrete selective, while voxels farther away in regions like temporoparietal junction (TPJ) tend to be abstract selective. These results replicate previous fMRI studies (Martin, 2016; Saxe and Kanwisher, 2003) mapping concrete and abstract concept representations across cortex.

**Figure 5.**
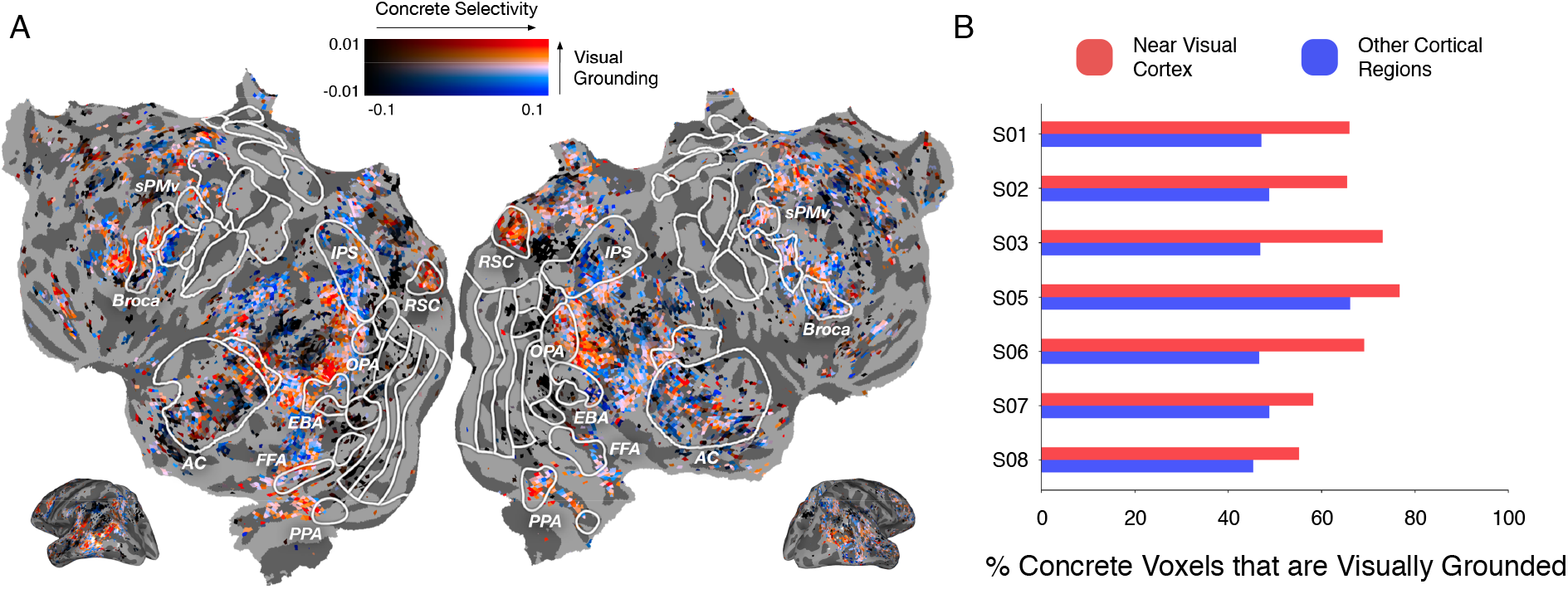
Representational format of concrete concepts across cortex. A *concrete selectivity score* was computed for each voxel as the projection of its encoding weights onto the vector of concreteness scores for each word. A *visual grounding score* was computed for each voxel as the difference in model performance between a visually grounded encoding model (*b* = -1) and a fully linguistic encoding model. **(A)** A cortical flatmap showing the concrete selectivity score and visual grounding score for each voxel in subject UT-S-02. Each semantic system voxel was assigned a brightness based on its concrete selectivity score and a color based on its visual grounding score. Concrete selective voxels were colored red if they are better modeled by the visually grounded space and blue if they are better modeled by the linguistic space. Abstract selective voxels were colored black. See **Figure S2** for similar maps for other subjects and visually grounded embedding spaces. Concrete selective voxels near the visual system are better modeled by the visually grounded space, while concrete selective voxels near somatosensory and motor systems are better modeled by the linguistic space. **(B)** The fraction of concrete selective voxels that are visually grounded was plotted near visual cortex, and in other cortical regions. For each subject, the fraction of concrete selective voxels that are visually grounded is higher near visual cortex than in other cortical regions.

Consistent with our previous results, we found that concrete selective voxels near visual cortex tend to be best modeled by the visually grounded space. Conversely, we found that concrete selective voxels in inferior parietal cortex and intraparietal sulcus (IPS) tend to be better modeled by the linguistic space than the visually grounded space. Based on their proximity to functional regions involved in somatosensory and motor processing, we predict that these parietal voxels represent concrete concepts using tactile features such as affordances (Barsalou, 2008; Binder and Desai, 2011), which may happen to be more aligned with the linguistic embedding space than the visual embedding space. The linguistic space also outperformed the visually grounded space in many inferior temporal voxels. While these regions are located near visual cortex, previous studies have suggested that they contain multimodal representations of object shape that combine visual and tactile information (Amedi et al., 2001). Notably, this visualization shows that concrete concepts are not invariably represented across cortex in a format that reflects visual information.

To quantify these results, we partitioned the set of semantic voxels with positive concrete selectivity scores into those located within 15mm of vision ROIs, and those located in other cortical regions. For each subset of concrete selective voxels, we computed the fraction with a positive visual grounding score (**Figure 5B**). Across subjects, 68 percent of concrete selective voxels near visual cortex were visually grounded, while only 49 percent of concrete selective voxels in other cortical regions were visually grounded. The fraction of concrete selective voxels that are visually grounded was significantly higher near visual cortex than in other cortical regions (*p* < 10^−3^, paired *t*-test; see **Methods**).

Together these results are consistent with the prediction that concrete concepts are represented near each perceptual system in a format that specifically reflects the corresponding modality. In particular, voxels near somatosensory and motor systems represent concrete concepts in a format that is *not* aligned with visual similarity, showing that concrete concepts are not invariably represented by visual information across cortex. However, because we do not explicitly model representations from non-visual perceptual systems, our results neither support nor challenge the existence of multimodal representations. While we have shown that certain concrete concept representations do not reflect visual information, it is possible that many voxels considered visually grounded in this study—particularly those farther from visual cortex (Binder and Desai, 2011)—may also reflect representations from other perceptual systems.

## Discussion

Most people learn about the world through both vision and language. This study characterized how these two sources of information are combined in the semantic system by modeling cortical concept representations evoked by narrative stories. We first operationalized visual and linguistic information as different embedding spaces, and then created a spectrum of semantic embedding spaces to model different possibilities for how visual and linguistic information are combined. Comparing encoding model performance between different semantic embedding spaces, we found that cortical regions near the visual system represent concepts using some amount of visual information, while cortical regions near the language system represent concepts using mostly linguistic information. Focusing on regions near visual cortex, we next demonstrated that most concepts are best modeled by a combination of visual and linguistic information, with more concrete concepts containing more visual information. Notably, however, we found that even many abstract concepts contain some amount of visual information from linguistically associated concrete concepts. Finally, we found that the visual grounding of concrete concepts—which tend to be experienced through multiple perceptual modalities—is localized near visual cortex, suggesting that semantic representations near each perceptual system specifically reflect how information is represented in the corresponding modality.

To facilitate future work in this area, we are sharing the semantic embedding spectrum and code used to generate it (https://github.com/jerryptang/grounded-embedding-spaces). Further, we plan to shortly release the entire fMRI dataset that was used in this study, which we hope will enable many future experiments since responses to natural language stimuli are highly reusable for asking many different scientific questions.

While we found consistent and statistically significant differences between semantic encoding models, these differences are numerically small. This is likely a consequence of the regression approach used to estimate the encoding models. In a regularized, ridge regression-based encoding model, weights are estimated to maximize the likelihood of the brain responses given the stimulus, under a prior that similar words in the embedding space should have similar weights (Nunez-Elizalde et al., 2019). However, as the amount of training data increases, the model can learn accurate weights from the data alone, decreasing the relative impact of the embedding space prior. Consequently, while our large fMRI dataset increases our confidence in the differences between embedding spaces, it also leads these differences to be numerically small.

Another potential issue is that the observed effects may not generalize beyond the narrative stories used to train and evaluate our encoding models. This issue of generalizability affects all fMRI experiments (Westfall et al., 2016). However, our study mitigates this issue to a large degree by using a very large set of natural language stimuli (5.37 hours or 55,144 total words) that span a broad space of semantic concepts, and an encoding framework in which we explicitly evaluate generalization performance of our models on multiple test stories. While issues of generalization can never be completely eliminated, our approach reduces this problem greatly compared to standard approaches in the field.

Our analyses are also bounded by our computational models of visual and linguistic representations. While our exploratory analyses (**Figure 2**) show that the visual and linguistic embedding spaces capture different notions of similarity, the embedding spaces are inherently imperfect models of visual and linguistic processing. Consequently, our results may be confounded by biases in the embedding spaces. For instance, we identified many voxels that are best modeled by semantic embedding spaces that solely contain linguistic information and concluded that these voxels represent concepts in a format that reflects linguistic representations (**Figure 3**) or representations from non-visual perceptual systems (**Figure 5**). However, we may also observe these results if the voxels contain visually grounded representations of concepts that are poorly modeled by the visual embedding space. This issue affects all model comparison experiments (Anderson et al., 2019). Our study attempts to mitigate this issue by using state-of-the-art computational models of visual and linguistic information. The analyses introduced in this study are applicable to all models that can be expressed as word embedding spaces, and can thus be used to test future models of visual and linguistic processing.

Finally, this study modeled semantic representations as combinations of visual and linguistic representations. However, there are many other sources through which humans acquire conceptual knowledge, such as somatosensation and emotion. We expect that some cortical regions that appear to reflect visual or linguistic representations may actually be best aligned with concept representations in these other modalities (**Figure 5**). Furthermore, other cortical regions may contain multimodal representations that combine information from multiple perceptual modalities (Binder and Desai, 2011). An important direction for future work is developing computational models for these other sources of information and using them to create increasingly detailed models of the semantic system.

## Methods

### MRI Data Collection

MRI data were collected on a 3T Siemens Skyra scanner at the UT Austin Biomedical Imaging Center using a 64-channel Siemens volume coil. Functional scans were collected using a gradient echo EPI sequence with repetition time (TR) = 2.00 s, echo time (TE) = 30.8 ms, flip angle = 71°, multi-band factor (simultaneous multi-slice) = 2, voxel size = 2.6mm x 2.6mm x 2.6mm (slice thickness = 2.6mm), matrix size = (84, 84), and field of view = 220 mm.

Anatomical data for all subjects except UT-S-02 were collected using a T1-weighted multi-echo MP-RAGE sequence on the same 3T scanner with voxel size = 1mm x 1mm x 1mm following the Freesurfer morphometry protocol. Anatomical data for subject UT-S-02 were collected on a 3T Siemens TIM Trio scanner at the UC Berkeley Brain Imaging Center using a 32-channel Siemens volume coil using the same sequence.

### Subjects

Data were collected from three female and four male human subjects: UT-S-01 (female, age 24), UT-S-02 (author A.G.H., male, age 34), UT-S-03 (male, age 22), UT-S-05 (female, age 23), UT-S-06 (author A.L., female, age 23), UT-S-07 (male, age 25), and UT-S-08 (male, age 24). All subjects were healthy and had normal hearing, and normal or corrected-to-normal vision. The experimental protocol was approved by the Institutional Review Board at the University of Texas at Austin. Written informed consent was obtained from all subjects. To stabilize head motion during scanning sessions participants wore a personalized head case that precisely fit the shape of each participant’s head (https://caseforge.co/).

### Natural Language Stimuli

The model estimation and evaluation data set consisted of 25 10-15 min stories taken from *The Moth Radio Hour*. In each story, a single speaker tells an autobiographical story without reading from a prepared speech. Each story was played during one scan with a buffer of 10 seconds of silence before and after the story. Data collection was broken up into 6 different scanning sessions, with the first session consisting of the anatomical scan and localizers, and each subsequent session consisting of 5 or 6 stories. A separate repeated test data set consisted of one 10 min story, also taken from *The Moth Radio Hour*. This story was played five times for each subject (once during each story scanning session), and the five sets of responses were averaged.

Stories were played over Sensimetrics S14 in-ear piezoelectric headphones. The audio for each story was filtered to correct for frequency response and phase errors induced by the headphones using calibration data provided by Sensimetrics and custom python code (https://github.com/alexhuth/sensimetrics_filter). All stimuli were played at 44.1 kHz using the pygame library in Python.

### fMRI Data Preprocessing

All functional data were motion corrected using the FMRIB Linear Image Registration Tool (FLIRT) from FSL 5.0. FLIRT was used to align all data to a template that was made from the average of all functional runs in the first story session for each subject. These automatic alignments were manually checked for accuracy. Low frequency voxel response drift was identified using a 2^nd^ order Savitzky-Golay filter with a 120 second window and then subtracted from the signal. To avoid onset artifacts and poor detrending performance near each end of the scan, responses were trimmed by removing 20 seconds (10 volumes) at the beginning and end of each scan, which removed the 10-second silent period and the first and last 10 seconds of each story. The mean response for each voxel was subtracted and the remaining response was scaled to have unit variance.

### Flatmap Construction

Cortical surface meshes were generated from the T1-weighted anatomical scans using FreeSurfer software (Dale et al., 1999). Before surface reconstruction, anatomical surface segmentations were hand-checked and corrected. Blender was used to remove the corpus callosum and make relaxation cuts for flattening. Functional images were aligned to the cortical surface using boundary based registration (BBR) implemented in FSL. These alignments were manually checked for accuracy and adjustments were made as necessary.

Flat maps were created by projecting the values for each voxel onto the cortical surface using the “nearest” scheme in pycortex software (Gao et al., 2015). This projection finds the location of each pixel in the flat map in 3D space and assigns that pixel the associated value.

### Stimulus Preprocessing

Each story was manually transcribed by one listener. Certain sounds (for example, laughter and breathing) were also marked to improve the accuracy of the automated alignment. The audio of each story was then downsampled to 11kHz and the Penn Phonetics Lab Forced Aligner (P2FA) (Yuan and Liberman, 2008) was used to automatically align the audio to the transcript. Praat (Boersma and Weenink, 2014) was then used to check and correct each aligned transcript manually.

### Localizers

Known regions of interest (ROIs) were localized separately in each subject. Three different tasks were used to define ROIs; a visual category localizer, an auditory cortex localizer, and a motor localizer.

Visual category localizer data were collected in six 4.5 minute scans consisting of 16 blocks of 16 seconds each. During each block 20 images of either places, faces, bodies, household objects, or spatially scrambled objects were displayed. Subjects were asked to pay attention to the same image being presented twice in a row. The cortical ROIs defined with this localizer were the fusiform face area (FFA), parahippocampal place area (PPA), occipital place area (OPA), retrosplenial cortex (RSC), and extrastriate body area (EBA).

Motor localizer data were collected in two identical 10 minute scans. The subject was cued to perform six different tasks in a random order in 20 second blocks. The cues were ‘hand’, ‘foot’, ‘mouth’, ‘speak’, saccade, and ‘rest’ presented as a word at the center of the screen, except for the saccade cue which was presented as an array of dots. For the ‘hand’ cue, subjects were instructed to make small finger-drumming movements for the entirety of the cue display. For the ‘foot’ cue, subjects were instructed to make small foot and toe movements. For the ‘mouth’ cue, subjects were instructed to make small vocalizations that were nonsense syllables such as *balabalabala*. For the ‘speak’ cue, subjects were instructed to self-generate a narrative without vocalization. For the saccade cue, subjects were instructed to make frequent saccades across the display screen for the duration of the task.

Weight maps for the motor areas were used to define primary motor and somatosensory areas for the hands, feet, and mouth; supplemental motor areas for the hands and feet, secondary somatosensory areas for the hands, feet, and mouth, and the ventral premotor hand area. The weight map for the saccade responses was used to define the frontal eye fields and intraparietal sulcus visual areas. The weight map for speech was used to define Broca’s area and the superior ventral premotor (sPMv) speech area (Chang et al., 2011).

Auditory cortex localizer data were collected in one 10 minute scan. The subject listened to 10 repeats of a 1-minute auditory stimulus containing 20 seconds of music (Arcade Fire), speech (Ira Glass, *This American Life*), and natural sound (a babbling brook). To determine whether a voxel was responsive to auditory stimulus, the repeatability of the voxel response across the 10 repeats was calculated using an *F-*statistic. This map was used to define the auditory cortex (AC).

### Visual and Linguistic Embedding Spaces

We constructed a linguistic embedding space based on word co-occurrence statistics in a large corpus of text (same as de Heer et al., 2017; Deniz et al., 2019; Huth et al., 2016). First, we constructed a 10,470-word lexicon from the union of the set of all words appearing in the first 2 story sessions and the 10,000 most common words in the large text corpus. We then selected 985 basis words from Wikipedia’s *List of 1000 Basic Words* (contrary to the title, this list contained only 985 unique words at the time it was accessed). This basis set was selected because it consists of common words that span a very broad range of topics. The text corpus used to construct this feature space includes the transcripts of 13 *Moth* stories (including 10 used as stimuli in this experiment), 604 popular books, 2,405,569 Wikipedia pages, and 36,333,459 user comments scraped from reddit.com. In total, the 10,470 words in our lexicon appeared 1,548,774,960 times in this corpus. Next, we constructed a word co-occurrence matrix, *L*, with 985 rows and 10,470 columns. Iterating through the text corpus, we added 1 to *L*_*i,j*_ each time word *j* appeared within 15 words of basis word *i*. A window size of 15 was selected to be large enough to suppress syntactic effects (that is, word order) but no larger. Once the word co-occurrence matrix was complete, we log-transformed the counts, replacing *L*_*i,j*_ with log(1 + *L*_*i,j*_). Next, each row of *L* was *z*-scored to correct for differences in basis word frequency, and then each column of *L* was *z*-scored to correct for word frequency. Each column of *L* is now a 985-dimensional vector representing one word in the lexicon. We then filtered the columns of *L* for the 3,933 unique words that occur in the stimulus stories. The linguistic embedding space is summarized by the covariance matrix Σ_*L*_ = *L*^T^*L*, where (Σ_*L*_)_*i,j*_ captures the degree of linguistic similarity between words *i* and *j*.

We constructed a visual embedding space based on embeddings extracted using a convolutional neural network (CNN). First, we defined a set of potential visual words from the union of words appearing in the first 2 story sessions and words with a concreteness rating *ċ* greater than or equal to 4.6 out of 5 in the Brysbaert Concreteness Ratings dataset (Brysbaert et al., 2014). We manually assigned each potential visual word the WordNet (Miller, 1995) synset that best corresponds to its linguistic meaning, which was inferred from the word’s 10 nearest neighbors in the linguistic embedding space Σ_*L*_. We then identified 720 visual words with ImageNet (Deng et al., 2009) entries corresponding to their assigned WordNet synsets. Of the 720 visual words, 394 were contained in the stimulus vocabulary. The 3,539 words in our stimulus vocabulary without corresponding ImageNet entries were considered non-visual. For each visual word, 100 images were randomly sampled from its ImageNet entry. 4,096-dimensional CNN embeddings were extracted for each image using the fc1 layer of a pretrained VGG16 (Simonyan and Zisserman, 2015) CNN implemented in Keras (Chollet and Others, 2015). We chose the feature extraction layer by fitting language encoding models (described below) induced by each layer of VGG16 on a single test subject (UT-S-02); fc1 attained the highest prediction performance across cortex. We obtained a CNN embedding for each visual word by averaging the extracted features across the 100 sampled images. The CNN embeddings were stored as columns in a matrix *C* with 4,096 rows and 720 columns.

We developed a perceptual propagation method to construct a matrix *V* of visual embeddings for both visual and non-visual words. We defined the linguistic submatrix *L*_*v*_ with 985 rows and 720 columns as the linguistic embeddings of the visual words. We then fit a linear model θ as *L*^T^ = θ*L*_*v*_^T^ to reconstruct each word’s linguistic embedding as a linear combination of the linguistic embeddings of visual words. For each word *w*, row θ_*w*_ contains 720 weights, which capture the degree to which each visual word contributes to the linguistic meaning of *w*. The matrix *V* of visual embeddings was then estimated by *V*^T^ = θ*C*^T^. *V* represents non-visual words as linear combinations of the CNN embeddings of associated visual words. *V* additionally combines each visual word’s CNN embedding with CNN embeddings of associated visual words, which smooths the visual embedding space (Collell et al., 2017). Finally, each column of *V*, which corresponds to the visual embedding of a word, was *z*-scored. The visual embedding space is summarized by the covariance matrix Σ_*V*_ = *V*^T^*V*, where (Σ_*V*_)_*i,j*_ captures the degree of visual similarity between words *i* and *j*.

We fit the perceptual propagation model θ using Tikhonov regression with prior covariance matrix Ω and regularization constant λ. We chose λ as the smallest value for which the first eigenvalue of the visual embedding space Σ_*V*_ was approximately equal to that of the linguistic space Σ_*L*_, in an effort to keep the smoothness of the visual embedding space as similar as possible to the linguistic embedding space. We tested two different prior covariance matrices; a spherical prior Ω_*I*_ that corresponds to ridge regression, and a CNN prior Ω_*C*_ = *C*^T^*C* which enforces that visual words with similar CNN embeddings have similar weights in θ. We found that for non-visual words, the associated visual words obtained under the spherical prior were more semantically diverse, while the associated visual words obtained under the CNN prior were more visually coherent. For example, the top associated words for “education” under the spherical prior were “school”, “college”, “university”, “student”, and “conservative”, while the top associated words under the CNN prior were “instructor”, “teacher”, “grade”, “student”, and “classroom” (which all depict a classroom setting). As the two priors capture different types of information, our perceptual propagation model θ was obtained by averaging the models θ_*I*_ and θ_*C*_.

### Concreteness Scores

We quantified the concreteness of each stimulus word using scores derived from the separate Brysbaert Concreteness Ratings dataset. The Brysbaert dataset contains human ratings *ċ* of the extent to which each word can be experienced through sensation. The concreteness ratings range from 1 (very abstract) to 5 (very concrete). We scaled the ratings between 0 (very abstract) and 1 (very concrete) by subtracting 1 and dividing by the range 4, and then squared the resulting values to obtain concreteness scores *c*. To interpolate concreteness scores for stimulus words that were not included in the Brysbaert dataset, each word *w* was assigned the max of its own concreteness score *c*_*w*_ (where *c*_*w*_ = 0 if *w* is not contained in the Brysbaert dataset) and the mean concreteness score of its 15 closest linguistic neighbors. Each word’s concreteness score *c*_*w*_ was thus given as 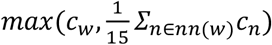, where the nearest neighbors function *nn*(*w*) gives the 15 closest words (where similarity is defined under Σ_*L*_) to *w* in the Brysbaert dataset.

### Visualizing Embedding Space Structure

We used PCA to visualize the structure of the visual and linguistic embedding spaces. For each space, we applied PCA to the embeddings of the 394 visual words that occur in the stimulus stories, and projected each word’s embedding onto the first two PCs. The first two PCs of the visual space account for 24.5% of the variance, and the first two PCs of the linguistic space account for 22.9% of the variance. For each embedding space, we plotted the two-dimensional projection of each visual word.

To highlight how notions of similarity differ between the visual and linguistic spaces, we identified 3 broad semantic categories; *people, clothes*, and *places*. For each category, we hand-selected 10 representative words prior to visualization, and colored the convex hull of the representative words in the two-dimensional visualization of each embedding space.

### Quantifying Word-level Differences in Embedding Spaces

For each word *w*, we defined a visual similarity vector (Σ_*V*_)_*w*_ containing its visual similarities with every other word, and a linguistic similarity vector (Σ_*L*_)_*w*_ containing its linguistic similarities with every other word. We computed a modality alignment score for each word as the linear correlation between its visual and linguistic similarity vectors. Words with high modality alignment scores are represented similarly in the visual and linguistic embedding spaces, while words with low modality alignment scores are represented differently in the visual and linguistic embedding spaces.

Across stimulus words, modality alignment scores *m* were anticorrelated with concreteness scores *c* (linear correlation *r* = -0.26). The linear least squares regression line between concreteness scores and modality alignment scores is *m* = -0.13*c* + 0.76.

### Semantic Embedding Spectrum

We created semantic embeddings *S*_*w*_ for each word *w* by concatenating its visual embedding *V*_*w*_ and its linguistic embedding *L*_*w*_. Each word was assigned a modality weight α_*w*_ between 0 and 1 to model the relative contributions of its visual and linguistic representations to its semantic representation. Prior to concatenation *V*_*w*_ was scaled to unit norm and then multiplied by α_*w*_^1/2^ while *L*_*w*_ was scaled to unit norm and then multiplied by (1 - α_*w*_)^1/2^. When α_*w*_ is 1 the semantic embedding *S*_*w*_ will fully reflect the visual embedding, and when α_*w*_ is 0 the semantic embedding *S*_*w*_ will fully reflect the linguistic embedding. Semantic embedding spaces are summarized by the covariance matrices Σ_*S*_ = *S*^T^*S*. The semantic similarity (Σ_*S*_)_*i,j*_ between words *i* and *j* is an average of their visual similarity Σ_*V*_ weighted by α_*i*_^1/2^α_*j*_^1/2^ and their linguistic similarity Σ_*L*_ weighted by (1 - α_*i*_)^1/2^(1 - α_*j*_)^1/2^.

Each semantic embedding space is parameterized by a vector **α** containing the modality weight α_*w*_ for each word *w*. To constrain the infinitely large space of **α** vectors we modeled each word’s modality weight α_*w*_ as a monotonically increasing function α_concrete_(*c* ; *b*) = σ(σ^-1^(*c*) + *b*) of its concreteness score *c*_*w*_, where σ is the sigmoid function σ(x) = e^x^/(e^x^ + 1). The α_concrete_ model has a single bias parameter *b* that controls the total amount of visual information in each word’s semantic embedding. As *b* approaches negative infinity, α(*c*_*w*_) approaches 0 for all *c*_*w*_, causing Σ_*S*_ to approach Σ_*L*_. As *b* approaches infinity, α(*c*_*w*_) approaches 1 for all *c*_*w*_, causing Σ_*S*_ to approach Σ_*V*_.

For our analyses, we chose 5 values of *b* (−10, -1, 0, 1, 10), which induce semantic embedding spaces that smoothly interpolate between the linguistic space Σ_*L*_ and the visual space Σ_*V*_. This semantic embedding spectrum contains a fully linguistic embedding space (*b* = -10) and a range of visually grounded embedding spaces (*b* = -1, 0, 1, 10)

### Voxelwise Encoding Models

fMRI encoding models are estimated on a set of training stories *S*_train_ and evaluated on a set of test stories *S*_test_. In model estimation, a response matrix *Y*_train_ is constructed by concatenating the fMRI responses to stories in *S*_train_. To construct the stimulus matrix *X*_train_, each word in *S*_train_ is first represented by a one-hot indicator vector corresponding to its identity in the 3,933-word stimulus vocabulary. The resulting binary matrix is then downsampled to the MR acquisition times using a 3-lobe Lanczos filter, yielding a *t*-by-3,933 dimensional word matrix *W*_train_, where *t* is the number of fMRI images in *Y*_train_. The word matrix *W*_train_ is then projected onto a feature matrix *P* which contains a *p*-dimensional embedding for each word, yielding the *t*-by-*p* dimensional stimulus matrix *X*_train_. Each feature channel of *X*_train_ is *z*-scored to match the features to the fMRI responses, which are *z*-scored within each story.

A linearized finite impulse response (FIR) model is fit to every cortical voxel in each subject’s brain. A separate linear temporal filter with four delays (1, 2, 3, and 4 time points) is fit for each of the *p* stimulus features, yielding a total of 4*p* features. This is accomplished by concatenating feature vectors that have been delayed by 1, 2, 3, and 4 time points (2, 4, 6, and 8 s). Taking the dot product of this concatenated feature space with a set of linear weights is functionally equivalent to convolving the original stimulus vectors with linear temporal kernels that have non-zero entries for 1-, 2-, 3-, and 4-time-point delays.

The 4*p* weights for each voxel are estimated from *X*_train_ and *Y*_train_ using L2-regularized linear regression (also known as ridge regression). The regression procedure has a single free parameter which controls the degree of regularization. This regularization coefficient is found for each voxel by repeating a regression and cross-validation procedure 50 times. In each iteration, approximately a fifth of the time points (*t* / 200 blocks of 40 consecutive time points each) are removed from the training data set and reserved for validation. Then the model weights are estimated on the remaining time points for each of 15 possible regularization coefficients (log spaced between 10 and 10,000). These weights are used to predict responses for the reserved time points, and prediction performance is computed between the predicted and actual responses. For each voxel, the regularization coefficient is chosen as the value that led to the best performance, averaged across bootstraps, on the reserved time points. For models where the sizes of the responses should be preserved (word-rate encoding models; described below), the regularization coefficient was optimized using *R*^2^ as the performance metric. For models where the sizes of the predicted responses do not matter (semantic encoding models; described below), the regularization coefficient was optimized using linear correlation as the performance metric.

The regression procedure produces a set of estimated feature weights *β*^*P*^, with columns corresponding to the 4*p* weights for each voxel. To evaluate a voxel-wise model, *β*^*P*^ is used to predict brain responses to stories in a test dataset *S*_test_ that were not used for model estimation. For each story *s* in *S*_test_, a stimulus matrix *X*_*s*_ and a response matrix *Y*_*s*_ are constructed using the procedure described above for constructing *X*_train_ and *Y*_train_. Each feature channel of *X*_*s*_ is normalized using the mean and standard deviation of the corresponding channel in *X*_train_. For each voxel, prediction performance on each test story is estimated as the linear correlation between predicted and actual responses over the time points in the story. Overall prediction performance on *S*_test_ is obtained by averaging the voxel’s prediction performance across the stories in *S*_test_.

### Encoding Model Estimation

Before fitting semantic encoding models, we first fit a word-rate encoding model for each subject to remove variance in the response data that could be explained by low-level auditory features. The word-rate model represents stimulus words with a 3,933-by-1 dimensional matrix of ones *P*_*WR*_. We estimated word-rate weights *β*_*WR*_ using all 5 story sessions as the training set *S*_*train*_. L2 regularization coefficients were chosen by maximizing *R*^2^ in the cross-validation procedure. For each of the 25 stimulus stories and the repeated test story, we predicted brain responses *Y*_*WR*_ = *Xβ*_*WR*_ using the word rate model. The word-rate predictions *Y*_*WR*_ were subtracted from the actual brain responses *Y*, which were then *z*-scored to produce word-rate corrected brain responses. Semantic encoding models were then fit to the word-rate corrected brain responses.

To fit a semantic encoding model with embedding space prior Σ, stimulus words were represented by embedding features *P* = Σ^1/2^. Previous work shows that performing ridge regression on the stimulus matrix *X* = *W*Σ^1/2^ is equivalent to performing Tikhonov regression on the word matrix *W* using Σ as the prior covariance (Nunez-Elizalde et al., 2019). L2 regularization coefficients were chosen by maximizing linear correlation in the cross-validation procedure. This procedure for solving Tikhonov regression yields a set of weights *β*^*P*^ on embedding features *P*. To represent the encoding model as weights on individual words, rather than weights on embedding features, we left-multiplied the feature space weights *β*^*P*^ by the delayed embedding features to obtain word-space weights *β*^*W*^ = (*I*_4_ ⊗ Σ^1/2^)*β*^*P*^. Each column of the weight matrix *β*^*W*^ contains a set of 15,732 estimated weights for a corresponding voxel. These weights predict how each of the 3,933 words in the stimulus vocabulary influences the BOLD responses in that voxel at each of the four temporal delays. When estimating the selectivity of each voxel for each word (**Figure 5**), we removed temporal information by averaging across the four delays for each word. Each voxel is then represented by a set of 3,933 averaged weights which predict how each word in the stimulus vocabulary influences the BOLD responses in that voxel.

To compare model performance under different embedding space priors (**Figure 3**), we estimated and evaluated encoding models using a bootstrap procedure across story sessions. For each of the 5 story sessions, we held out the chosen session as *S*_test_ and estimated encoding models using the remaining 4 story sessions as *S*_train_. We then computed prediction performance of the estimated models on each story in *S*_test_. Repeating this process for each story session yielded prediction performance on all 25 stimulus stories. Aggregate performance was obtained by averaging performance across the 25 stories. As the stimulus stories vary in semantic content and imageability, maximizing the number of evaluation stories was desirable for identifying the embedding space that best models each voxel. Because this session bootstrap procedure evaluated encoding models on single repetitions of many stories rather than many repetitions of a single story (de Heer et al., 2017; Huth et al., 2016; Jain and Huth, 2018), our reported prediction performance values were lower than previously reported results due to the lower signal-to-noise ratio of single repetition response data.

A downside to the story session bootstrap procedure is that the 5 story sessions produce 5 separate encoding models. As the encoding models were not estimated using independent data, their weights cannot be meaningfully combined. Furthermore, the story session bootstrap procedure is computationally intensive. For analyses estimating voxel selectivity from encoding model weights (**Figure 5**) and analyses that compare a large number of encoding models (**Figure 4**), we instead split the story sessions into explicit train and test sessions. This procedure produces a single set of encoding model weights. The number of training and test sessions used depends on the nature of each analysis, as described below.

All model fitting and analysis was performed using custom software written in Python, making heavy use of NumPy (Oliphant, 2006), SciPy (Jones et al., 2001), and pycortex (Gao et al., 2015).

### Semantic System Voxels

Semantic system voxels were defined as voxels that were significantly predicted by any space in the semantic embedding spectrum. We tested for significance using a permutation test on the repeated test story *S*_reptest_. The embedding spectrum performance for each voxel was defined as the maximum linear correlation *r* between the true response time course and the predicted response time course under each semantic embedding space. We then constructed a null distribution on embedding spectrum performance for each voxel by permuting the voxel’s true response time course. In each trial, we randomly resampled (with replacement) 10-TR blocks from the voxel’s true response time course. Resampling contiguous blocks preserves the auto-correlation structure of the voxel’s responses. We then computed null embedding spectrum performance as the maximum linear correlation *r* between the permuted response time course and the predicted response time course under each semantic embedding space. Repeating this process for 10,000 trials provided a null distribution of embedding spectrum performance for each voxel. Semantic system voxels were identified as voxels with an observed embedding spectrum performance that is significantly higher than its null distribution (q(FDR) < 0.05), correcting for multiple comparisons using the false discovery rate (Benjamini and Hochberg, 1995).

For encoding models estimated using the session bootstrap procedure (**Figures 3, 5**) we averaged across the 5 sets of encoding weights (corresponding to each bootstrap session) to predict responses to the repeated test story. This yielded 8,578 semantic system voxels in subject UT-S-01, 13,502 semantic system voxels in UT-S-02, 17,135 semantic system voxels in UT-S-03, 3,835 semantic system voxels in UT-S-05, 5,504 semantic system voxels in UT-S-06, 3,065 semantic system voxels in UT-S-07, and 1,321 semantic system voxels in UT-S-08.

For encoding models estimated using an explicit train-test split (**Figure 4**) we predicted responses to the repeated test story using the single set of encoding weights. This yielded 7,047 semantic system voxels in subject UT-S-01, 11,933 semantic system voxels in UT-S-02, 12,807 semantic system voxels in UT-S-03, 3,338 semantic system voxels in UT-S-05, 2,539 semantic system voxels in UT-S-06, 2,230 semantic system voxels in UT-S-07, and 807 semantic system voxels in UT-S-08.

### Linear Mixed-effects Modeling

A linear mixed-effects model (lme) was used to compare the performance of different spaces in the semantic embedding spectrum around vision and language ROIs. We identified vision (FFA, PPA, OPA, RSC, EBA) and language (AC, Broca, sPMv) ROIs in each subject using separate localizer data (described above). We used pycortex software (Gao et al., 2015) to identify semantic system voxels within 15mm of each ROI along the cortical surface. For each ROI, we first identified all vertices on the fiducial surface that fall within the ROI definition. We then computed the geodesic distance from each surface vertex to the closest vertex in the ROI. We defined ROI-adjacent vertices as vertices within 15mm of the ROI vertices. We finally used the “cortical” scheme in pycortex to select all voxels with centers within the cortical ribbon where the closest vertex is ROI-adjacent.

For each subject, the performance of each embedding space around an ROI was computed by averaging the prediction performance of the corresponding encoding model (estimated under the story session bootstrap encoding procedure) across semantic system voxels within 15mm of the ROI. We then computed a visual grounding score for each visually grounded embedding space as its performance improvement over the fully linguistic embedding space. Our linear mixed-effects model compared visual grounding score for each visually grounded embedding space (4 levels: *b* = -1, 0, 1, 10) and ROI type (2 levels: vision, language). The ROI ID nested within subject ID was the random effect. The lme test was run in R using the lme4 library (Bates et al., 2015). For post hoc tests, *p*-values were corrected for multiple comparisons using the false discovery rate.

For each ROI, we plotted the visual grounding score for each visually grounded embedding space. We then plotted mean visual grounding score across vision and language ROIs for each visually grounded embedding space. All values were averaged across 7 subjects. Error bars indicate standard error of the mean across 7 subjects.

### Modality Weight Permutation Test

We conducted a two-tailed permutation test to determine whether the amount of visual information in each word’s semantic representation around visual cortex is related to concreteness. We first identified the best α_concrete_ model around visual cortex (*b* = -1) by comparing encoding model performance on the repeated test set *S*_reptest_. We then fit semantic encoding models using the first 3 story sessions as *S*_train_ and the remaining 2 story sessions as *S*_test_. L2 regularization coefficients were chosen by maximizing linear correlation in the cross-validation procedure. Encoding model performance (linear correlation *r*) was averaged across semantic system voxels within 15mm of vision ROIs (FFA, PPA, OPA, RSC, EBA) along the cortical surface.

We next conducted 1,000 trials in which we permuted concreteness scores across words before computing modality weights under the α_concrete_ model (*b* = -1). In trial *t* of the permutation test, the modality weights across stimulus words were given by a vector **α**_*t*_ corresponding to a random permutation of the concreteness-derived modality weights **α**_concrete_. We then fit an encoding model under the semantic embedding space induced by **α**_*t*_ and averaged encoding model performance across the tested voxels. For each voxel, we reused the L2 regularization coefficient previously optimized for the α_concrete_ encoding model.

The 1,000 trials provide a permutation distribution of the encoding model performance. The permutation distribution was significantly lower than the observed performance of the α_concrete_ model when combined across subjects (q(FDR) < 10^−4^), and individually for five of seven subjects (q(FDR) < 10^−2^).

### Binary Modality Weight Model

The visually grounded parameterizations (*b* = -1, 0, 1, 10) of the α_concrete_ modality weight model predict that all abstract words contain some amount of visual information. To capture the alternative hypothesis that abstract words solely contain linguistic information, we defined an α_binary_ modality weight model parameterized by an abstractness cutoff *a*. Words with concreteness scores below the cutoff were considered purely abstract and represented solely by their linguistic embeddings, while words with concreteness scores above than the cutoff were represented by a combination of their visual and linguistic embeddings specified in the α_concrete_ model. Formally, α_binary_(*c*; *a, b*) is a piecewise function that outputs 0 if *c* is less than *a*, and α_concrete_(*c*; *b*) otherwise. To directly compare α_concrete_ and α_binary_, both models were parameterized by the best bias parameter for α_concrete_ around visual cortex (*b* = -1), which was determined by comparing encoding model performance on the repeated test set *S*_reptest_.

We compared the α_concrete_ model against the α_binary_ model for a range of abstractness cutoffs (*a* = 0.1, 0.2, 0.3, 0.4, 0.5, 0.6, 0.7, 0.8, 0.9, 1.0). For each modality weight model, we fit a semantic encoding model under the induced embedding space using the first 3 story sessions as *S*_train_ and the remaining 2 story sessions as *S*_test_. For both the α_concrete_ and α_binary_ encoding models, L2 regularization coefficients were chosen by maximizing linear correlation in the cross-validation procedure. Encoding model performance (linear correlation *r*) was averaged across semantic system voxels within 15mm of vision ROIs (FFA, PPA, OPA, RSC, EBA) along the cortical surface.

A linear mixed-effects model (lme) was used to compare the performance difference between each α_binary_ model and the α_concrete_ model (11 levels: *a* = 0.1, 0.2, 0.3, 0.4, 0.5, 0.6, 0.7, 0.8, 0.9, 1.0). The subject ID was the random effect. The lme test was run in R using the lme4 library (Bates et al., 2015). For post hoc tests, *p*-values were corrected for multiple comparisons using the false discovery rate. α_binary_ models with concrete cutoffs of 0.6, 0.8, 0.9, and 1.0 performed significantly worse than the α_concrete_ model (q(FDR) < 0.05).

### Visual Grounding of Concrete Selective Voxels

We defined a concrete selectivity score for each voxel to quantify the degree to which it responds to concrete words. We fit encoding models under the fully linguistic embedding space using all 5 story sessions as *S*_train_. The estimated encoding weights (averaged across delays) predict the degree to which each word influences BOLD responses in each voxel. We then projected a vector of concreteness scores for each word onto the encoding weights for each voxel. We divided each voxel’s score by the sum of its absolute weights on each word. Concrete selectivity scores range from -1 to 1; voxels that respond more to concrete words than abstract words will have positive concrete selectivity scores, while voxels that respond more to abstract words than concrete words will have negative concrete selectivity scores.

We defined a visual grounding score for each voxel to quantify the degree to which it represents concepts in a visually grounded format. We determined the best visually grounded parameterization of α_concrete_ across visual cortex (*b* = -1) by comparing encoding model performance on the repeated test set *S*_reptest_. The visual grounding score of each voxel was then defined as the difference in encoding model performance (estimated under the story session bootstrap procedure) between the visually grounded embedding space (*b* = -1) and the fully linguistic embedding space (*b* = -10). Visual grounding scores range from -1 to 1; voxels that represent concepts in a visually grounded format will have positive visual grounding scores, while voxels that represent concepts in a linguistic format will have negative visual grounding scores.

We defined concrete selective voxels as semantic system voxels with a positive concrete selectivity score. We tested whether concrete selective voxels are more visually grounded near visual cortex than in other cortical regions. We partitioned concrete selective voxels into those near visual cortex (within 15mm of visual ROIs) and those in other cortical regions. For each subset of concrete selective voxels, we computed the fraction that are visually grounded (visual grounding score > 0). Combined across subjects, 68 percent of concrete selective voxels near visual cortex were visually grounded, while 49 percent of concrete selective voxels in other cortical regions were visually grounded. We conducted a two-tailed paired t-test across subjects comparing the fraction of concrete selective voxels near visual cortex that are visually grounded to the fraction of concrete selective voxels in other cortical regions that are visually grounded. We found that concrete selective voxels were significantly more likely to be visually grounded near visual cortex than in other cortical regions (*p* < 0.01).

## Supplemental Figures

**Figure S1 (related to Figure 3).**
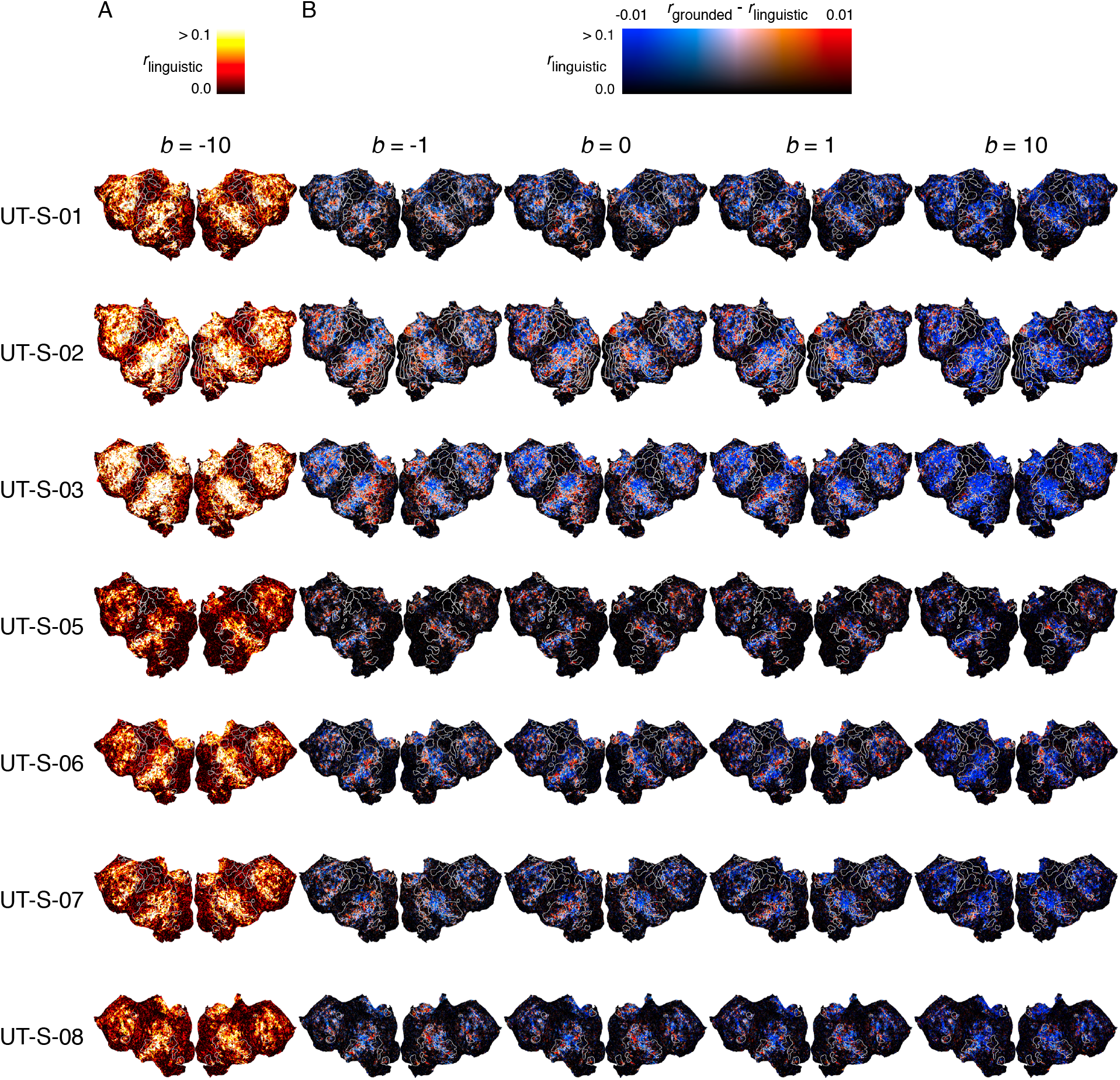
Encoding model performance across semantic embedding spaces. Encoding models were fit using each space in a semantic embedding spectrum ranging from fully linguistic to fully visual. Prediction performance for each voxel is measured by mean linear correlation *r* across 25 evaluation stories. **(A)** Cortical flatmaps show the prediction performance of the fully linguistic embedding space (*b* = -10) for each voxel in each subject. Well-predicted voxels appear yellow or white, and poorly predicted voxels appear black. **(B)** Cortical flatmaps show the difference in prediction performance between each visually grounded embedding space and the fully linguistic embedding space. Voxels that are better predicted by each visually grounded space are colored red, and voxels that are better predicted by the fully linguistic space are colored blue. The brightness of each voxel is given by the performance of the fully linguistic space.

**Figure S2 (related to Figure 5).**
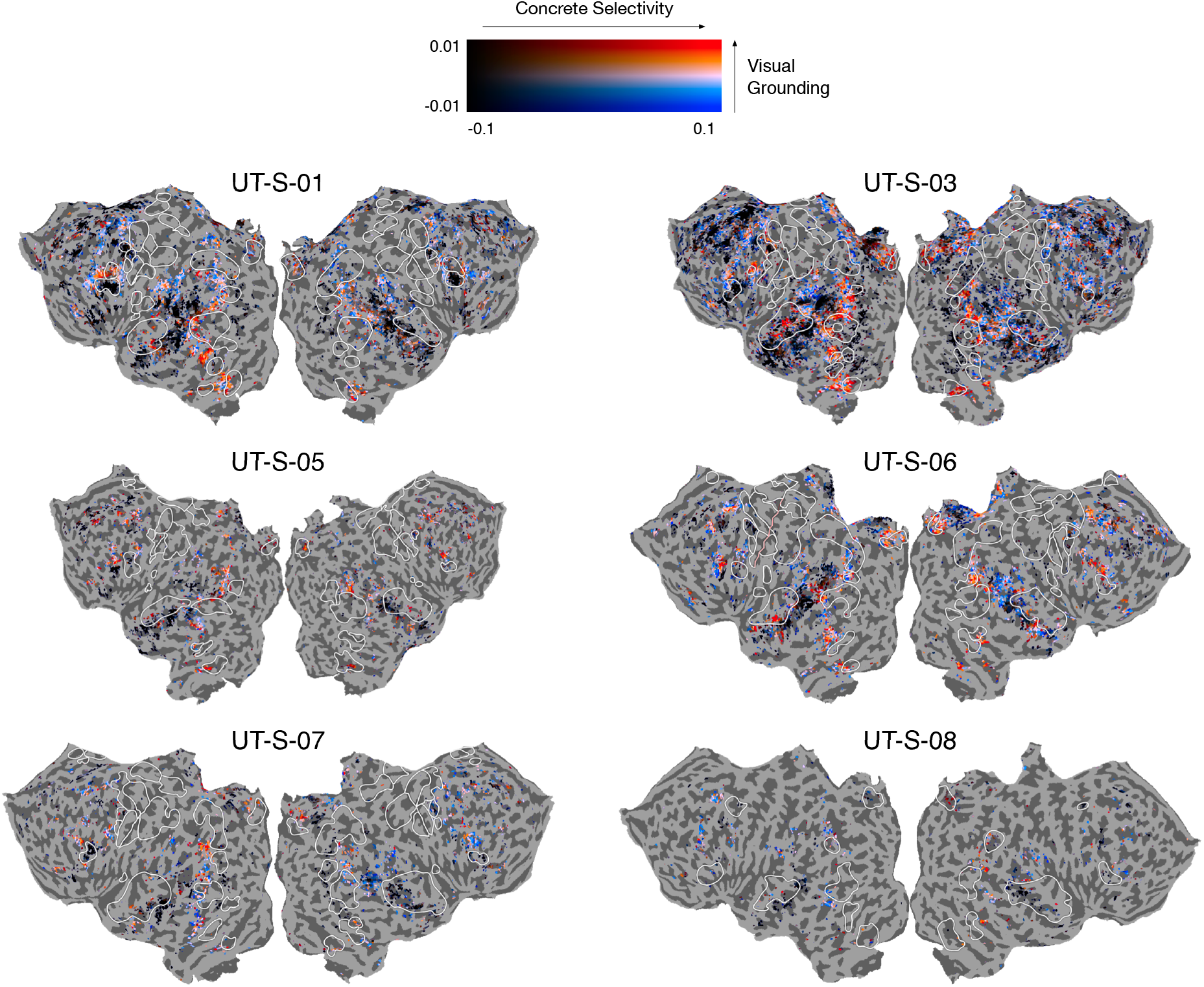
Representational format of concrete concepts across cortex. Similar to **Figure 5** in the main text, a *concrete selectivity score* was computed for each voxel as the projection of its encoding weights onto the vector of concreteness scores for each word, and a *visual grounding score* was computed for each voxel as the difference in model performance between a visually grounded encoding model (*b* = -1) and a fully linguistic encoding model. Cortical flatmaps show the concrete selectivity score and visual grounding score for each voxel in subjects UT-S-01, UT-S-03, UT-S-05, UT-S-06, UT-S-07, and UT-S-08. These maps show that across subjects, concrete selective voxels near the visual system are better modeled by the visually grounded space, while concrete selective voxels near somatosensory and motor systems are better modeled by the linguistic space.

## References

Amedi A, Malach R, Hendler T, Peled S, Zohary E. 2001. Visuo-haptic object-related activation in the ventral visual pathway. Nat Neurosci 4:324–330.

Anderson AJ, Binder JR, Fernandino L, Humphries CJ, Conant LL, Raizada RDS, Lin F, Lalor EC. 2019. An Integrated Neural Decoder of Linguistic and Experiential Meaning. J Neurosci 39:8969–8987.

Andrews M, Frank S, Vigliocco G. 2014. Reconciling embodied and distributional accounts of meaning in language. Top Cogn Sci 6:359–370.

Barsalou LW. 2008. Grounded cognition. Annu Rev Psychol 59:617–645.

Bates D, Mächler M, Bolker B, Walker S. 2015. Fitting Linear Mixed-Effects Models Using lme4. Journal of Statistical Software. doi:10.18637/jss.v067.i01

Benjamini Y, Hochberg Y. 1995. Controlling the False Discovery Rate: A Practical and Powerful Approach to Multiple Testing. J R Stat Soc Series B Stat Methodol 57:289–300.

Binder JR, Desai RH. 2011. The neurobiology of semantic memory. Trends Cogn Sci 15:527– 536.

Binder JR, Desai RH, Graves WW, Conant LL. 2009. Where is the semantic system? A critical review and meta-analysis of 120 functional neuroimaging studies. Cereb Cortex 19:2767– 2796.

Binder JR, Westbury CF, McKiernan KA, Possing ET, Medler DA. 2005. Distinct brain systems for processing concrete and abstract concepts. J Cogn Neurosci 17:905–917.

Boersma P, Weenink D. 2014. Praat: doing phonetics by computer.

Borghi AM, Binkofski F, Castelfranchi C, Cimatti F, Scorolli C, Tummolini L. 2017. The challenge of abstract concepts. Psychol Bull 143:263–292.

Bruni E, Tran N-K, Baroni M. 2014. Multimodal distributional semantics. J Artif Intell Res 49:1– 47.

Brysbaert M, Warriner AB, Kuperman V. 2014. Concreteness ratings for 40 thousand generally known English word lemmas. Behav Res Methods 46:904–911.

Cadieu CF, Hong H, Yamins DLK, Pinto N, Ardila D, Solomon EA, Majaj NJ, DiCarlo JJ. 2014. Deep neural networks rival the representation of primate IT cortex for core visual object recognition. PLoS Comput Biol 10:e1003963.

Chang EF, Edwards E, Nagarajan SS, Fogelson N, Dalal SS, Canolty RT, Kirsch HE, Barbaro NM, Knight RT. 2011. Cortical spatio-temporal dynamics underlying phonological target detection in humans. J Cogn Neurosci 23:1437–1446.

Chatfield K, Simonyan K, Vedaldi A, Zisserman A. 2014. Return of the Devil in the Details: Delving Deep into Convolutional Nets. arXiv [csCV].

Chollet F, Others. 2015. Keras. https://keras.io

Collell G, Zhang T, Moens M-F. 2017. Imagined visual representations as multimodal embeddingsThirty-First AAAI Conference on Artificial Intelligence.

Dale AM, Fischl B, Sereno MI. 1999. Cortical surface-based analysis. I. Segmentation and surface reconstruction. Neuroimage 9:179–194.

Deerwester S, Dumais ST, Furnas GW, Landauer TK, Harshman R. 1990. Indexing by latent semantic analysis. Journal of the American society for information science 41:391–407.

de Heer WA, Huth AG, Griffiths TL, Gallant JL, Theunissen FE. 2017. The hierarchical cortical organization of human speech processing. Journal of Neuroscience 37:6539–6557.

Deng J, Dong W, Socher R, Li L-J, Li K, Fei-Fei L. 2009. ImageNet: A large-scale hierarchical image database2009 IEEE Conference on Computer Vision and Pattern Recognition. IEEE. pp. 248–255.

Deniz F, Nunez-Elizalde AO, Huth AG, Gallant JL. 2019. The Representation of Semantic Information Across Human Cerebral Cortex During Listening Versus Reading Is Invariant to Stimulus Modality. J Neurosci 39:7722–7736.

Dove G. 2009. Beyond perceptual symbols: a call for representational pluralism. Cognition 110:412–431.

Eickenberg M, Gramfort A, Varoquaux G, Thirion B. 2017. Seeing it all: Convolutional network layers map the function of the human visual system. Neuroimage 152:184–194.

Gao JS, Huth AG, Lescroart MD, Gallant JL. 2015. Pycortex: an interactive surface visualizer for fMRI. Front Neuroinform 9:23.

Glenberg AM, Robertson DA. 2000. Symbol grounding and meaning: A comparison of high-dimensional and embodied theories of meaning. J Mem Lang 43:379–401.

Güçlü U, van Gerven MAJ. 2015. Deep neural networks reveal a gradient in the complexity of neural representations across the ventral stream. Journal of Neuroscience 35:10005– 10014.

Hamilton LS, Huth AG. 2018. The revolution will not be controlled: natural stimuli in speech neuroscience. Language, Cognition and Neuroscience 1–10.

Harnad S. 1990. The symbol grounding problem. Physica D.

Harpaintner M, Sim E-J, Trumpp NM, Ulrich M, Kiefer M. 2020. The grounding of abstract concepts in the motor and visual system: An fMRI study. Cortex 124:1–22.

Harpaintner M, Trumpp NM, Kiefer M. 2018. The Semantic Content of Abstract Concepts: A Property Listing Study of 296 Abstract Words. Front Psychol 9:1748.

Huth AG, de Heer WA, Griffiths TL, Theunissen FE, Gallant JL. 2016. Natural speech reveals the semantic maps that tile human cerebral cortex. Nature 532:453.

Jain S, Huth A. 2018. Incorporating Context into Language Encoding Models for fMRIAdvances in Neural Information Processing Systems. pp. 6629–6638.

Jones E, Oliphant T, Peterson P. 2001. SciPy: Open source scientific tools for Python.

Khaligh-Razavi S-M, Kriegeskorte N. 2014. Deep supervised, but not unsupervised, models may explain IT cortical representation. PLoS Comput Biol 10:e1003915.

Krizhevsky A, Sutskever I, Hinton GE. 2012. Imagenet classification with deep convolutional neural networksAdvances in Neural Information Processing Systems. pp. 1097–1105.

Lund K, Burgess C. 1996. Producing high-dimensional semantic spaces from lexical co-occurrence. Behav Res Methods Instrum Comput 28:203–208.

Lynott D, Connell L, Brysbaert M, Brand J, Carney J. 2020. The Lancaster Sensorimotor Norms: multidimensional measures of perceptual and action strength for 40,000 English words. Behav Res Methods 52:1271–1291.

Martin A. 2016. GRAPES—Grounding representations in action, perception, and emotion systems: How object properties and categories are represented in the human brain. Psychon Bull Rev 23:979–990.

Miller GA. 1995. WordNet: a lexical database for English. Commun ACM 38:39–41.

Mitchell TM, Shinkareva SV, Carlson A, Chang K-M, Malave VL, Mason RA, Just MA. 2008. Predicting human brain activity associated with the meanings of nouns. Science 320:1191– 1195.

Nunez-Elizalde AO, Huth AG, Gallant JL. 2019. Voxelwise encoding models with non-spherical multivariate normal priors. Neuroimage 197:482–492.

Oliphant TE. 2006. A guide to NumPy. Trelgol Publishing USA.

Paivio A. 1991. Dual coding theory: Retrospect and current status. Canadian Journal of Psychology/Revue canadienne de psychologie 45:255–287.

Pennington J, Socher R, Manning C. 2014. Glove: Global vectors for word representationProceedings of the 2014 Conference on Empirical Methods in Natural Language Processing (EMNLP). pp. 1532–1543.

Riordan B, Jones MN. 2011. Redundancy in perceptual and linguistic experience: Comparing feature-based and distributional models of semantic representation. Top Cogn Sci 3:303– 345.

Saxe R, Kanwisher N. 2003. People thinking about thinking peopleThe role of the temporo-parietal junction in “theory of mind.” NeuroImage. doi:10.1016/s1053-8119(03)00230-1

Sermanet P, Eigen D, Zhang X, Mathieu M, Fergus R, LeCun Y. 2013. OverFeat: Integrated Recognition, Localization and Detection using Convolutional Networks. arXiv [csCV].

Simonyan K, Zisserman A. 2015. Very Deep Convolutional Networks for Large-Scale Image RecognitionInternational Conference on Learning Representations.

Wehbe L, Murphy B, Talukdar P, Fyshe A, Ramdas A, Mitchell T. 2014. Simultaneously uncovering the patterns of brain regions involved in different story reading subprocesses. PLoS One 9:e112575.

Westfall J, Nichols TE, Yarkoni T. 2016. Fixing the stimulus-as-fixed-effect fallacy in task fMRI. Wellcome open research 1.

Yamins DLK, Hong H, Cadieu CF, Solomon EA, Seibert D, DiCarlo JJ. 2014. Performance-optimized hierarchical models predict neural responses in higher visual cortex. Proc Natl Acad Sci U S A 111:8619–8624.

Yuan J, Liberman M. 2008. Speaker identification on the SCOTUS corpus. J Acoust Soc Am 123:3878.

Zeiler MD, Fergus R. 2014. Visualizing and Understanding Convolutional NetworksComputer Vision – ECCV 2014. Springer International Publishing. pp. 818–833.

